# UBR-1 enzyme network regulates glutamate homeostasis to affect organismal behavior and developmental viability

**DOI:** 10.1101/2025.07.28.666006

**Authors:** Joseph S. Pak, Seamus Morrone, Karla J. Opperman, Mukul K. Midha, Charu Kapil, Neal D. Mathew, Damon T. Page, Ning Zheng, Robert L. Moritz, Brock Grill

## Abstract

Johanson-Blizzard Syndrome (JBS) is an autosomal recessive spectrum disorder associated with the UBR-1 ubiquitin ligase that features developmental delay including motor abnormalities. Here, we demonstrate that *C. elegans* UBR-1 regulates high-intensity locomotor behavior and developmental viability via both ubiquitin ligase and scaffolding mechanisms. Super-resolution imaging with CRISPR-engineered UBR-1 and genetic results demonstrated that UBR-1 is expressed and functions in the nervous system including in pre-motor interneurons. To decipher mechanisms of UBR-1 function, we deployed CRISPR-based proteomics using *C. elegans* which identified a cadre of glutamate metabolic enzymes physically associated with UBR-1 including GLN-3, GOT-2.2, GFAT-1 and GDH-1. Similar to UBR-1, all four glutamate enzymes are genetically linked to human developmental and neurological deficits. Proteomics, multi-gene interaction studies, and pharmacological findings indicated that UBR-1, GLN-3 and GOT-2.2 form a signaling axis that regulates glutamate homeostasis. Developmentally, UBR-1 is expressed in embryos and functions with GLN-3 to regulate viability. Overall, our results suggest UBR-1 is an enzyme hub in a GOT-2.2/UBR-1/GLN-3 axis that maintains glutamate homeostasis required for efficient locomotion and organismal viability. Given the prominent role of glutamate within and outside the nervous system, the UBR-1 glutamate homeostatic network we have identified could contribute to JBS etiology.

## Introduction

UBR1 is a RING-type E3 ligase that is conserved from yeast through multiple animal lineages. It is a member of the UBR-box ligases that regulate protein quality control through the N-end rule pathway ^1^7/28/25 1:51:00 PM. In humans, UBR1 is associated with a rare autosomal recessive genetic disorder called Johanson-Blizzard Syndrome (JBS) ^2–6^. JBS patients display a variable spectrum of developmental abnormalities that includes developmental delay, facial abnormalities, deafness, and cognitive impairment. While defective pancreas development has been extensively studied in JBS and *Ubr1* mutant mice, our understanding of UBR1 function and JBS pathophysiology in the nervous system remains relatively limited.

Studies with yeast and human cell-based models have provided important progress on understanding UBR1 functions, which include regulating apoptosis, mitochondrial quality control, and G-protein signaling ^7–11^. However, much less is known about the molecular and cellular roles of UBR1 *in vivo*. While JBS patients display neurodevelopmental abnormalities, we still have a relatively limited understanding of UBR1 function in the nervous system. *Ubr1* mutant mice exhibit decreased muscle mass, which could reflect problems with NMJ function ^2,12^. However, few other nervous system phenotypes have been observed in *Ubr1* mutant mice, which could be due to functional redundancy between UBR1 and its paralog UBR2. Consistent with this, mouse double knockouts for *Ubr1* and *Ubr2* display defects in neurogenesis, suggesting an important role in the nervous system ^13^. To date, embryonic lethality of mouse double knockouts has limited further organismal and neurological studies.

Here, we present our findings on UBR-1 using the invertebrate *C. elegans* as an organismal model for behavioral and developmental studies. Two key advantages of *C. elegans* are that it has a sole ortholog for mammalian UBR1 and UBR2, and *ubr-1* mutants are not lethal. As a result, we were able to show that *ubr-1* mutants have robust defects in high-intensity locomotor behavior and reduced organismal viability. Using unbiased CRISPR-based proteomics in *C. elegans*, we show that UBR-1 is a scaffold for a network of enzymes responsible for glutamate metabolism and signaling.

Glutamate is a primary, conserved neurotransmitter from *C. elegans* through mammals ^14–16^. Glutamate is also required for protein synthesis and is a precursor for the citric acid cycle. Balanced glutamate levels, or glutamate homeostasis at an organismal level, are important for nervous system health and function. For example, accumulation of excess glutamate in the nervous system results in increased glutamatergic transmission, neuronal overactivation, and excitotoxicity ^17,18^. Alterations in nervous system excitatory and inhibitory balance has also been implicated in a range of neurodevelopmental disorders ^19^. In the brains of vertebrates, astrocytes also play a role in glutamate homeostasis by taking up extracellular glutamate, and converting glutamate to glutamine which is shuttled back to presynaptic terminals ^20^. Finally, disrupted homeostasis of glutamate and glutamine is implicated in epilepsy, encephalopathy, and neurodegeneration ^21,22,20^.

At present, our understanding of how ubiquitin ligases functionally integrate with the glutamate enzymatic machinery remains relatively poorly understood. Prior studies showed that the RBX ubiquitin ligase forms a complex with a mitochondrial glutamate metabolic enzyme GDH1, and ubiquitin ligases such as the APC have been found to regulate glutamatergic ion channels in the nervous system ^23–26^. However, it remains unclear whether ubiquitin ligases influence glutamate homeostasis to shape nervous system function and organismal development. Here, we show that UBR-1 functions as an enzyme hub in a GOT-2.2/UBR-1/GLN-3 axis that regulates glutamate homeostasis to influence locomotor behavior and organismal viability.

## Results

### UBR-1 regulates high-intensity locomotor behavior in C. elegans

*C. elegans* UBR-1 is an ancient RING family ubiquitin ligase that is conserved from yeast through mammals (**Supplementary Fig 1**). Patients with JBS carry genetic variants in *UBR1*, display developmental delay, and miss motor development milestones ^3,4^. Prior studies in *C. elegans* have indicated that *ubr-1* mutants have defects in body bends during reversal on solid media ^27^. To more broadly evaluate how UBR-1 regulates locomotor behavior, we examined *ubr-1* mutants using high-intensity locomotor assays in liquid. Importantly, swimming is a much more vigorous and distinct form of *C. elegans* locomotion compared to more intermittent crawling behavior on solid media ^28,29^.

We began by testing a *ubr-1* mutant, *tm5996*, which is an out-of-frame deletion that leads to an early stop (H137fs*) generating a likely molecular null (**Fig 1A**). To initially evaluate locomotor behavior, we generated movies of wild-type animals and *ubr-1 (tm5996)* mutants swimming in buffer (**Fig 1B; Supplementary Movies 1&2**). Substantially reduced locomotion was clearly apparent in *ubr-1* mutants in these high intensity locomotor assays. In contrast, forward locomotor defects on solid media plates were not observed for *ubr-1* mutants (**Supplementary Fig 2; Supplementary Movies 3&4)**. We also observed an exaggerated lack of coordination in tail movements of *ubr-1* mutants while swimming (**Fig 1B**), consistent with prior results for reversal studies on solid media ^27^.

**Fig 1.**
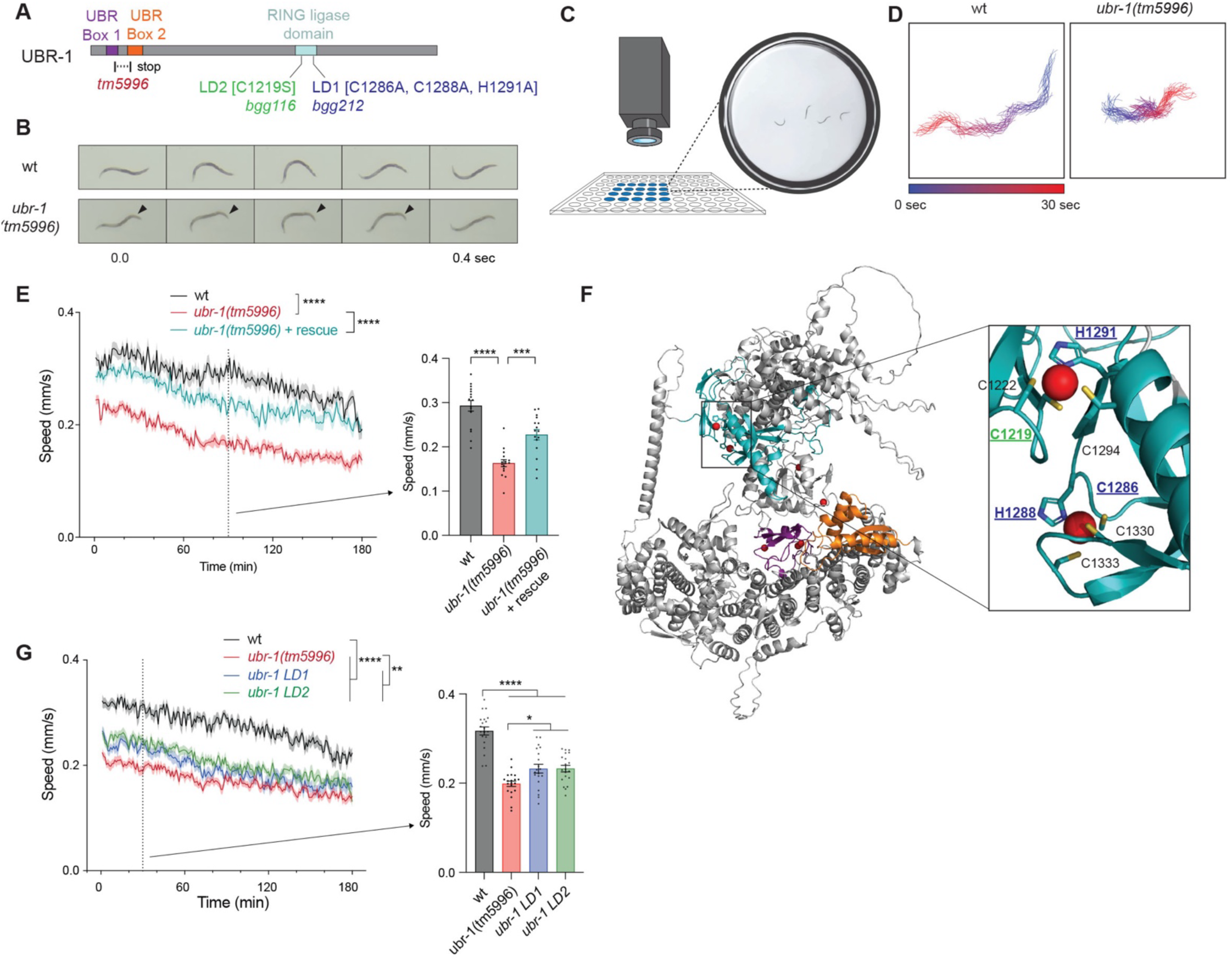
UBR-1 regulates high-intensity locomotor behavior in *C. elegans*. **A**. Schematic of UBR-1 and annotated protein domains showing deletion allele (*tm5996*) and CRISPR engineered RING inactivation alleles, *LD1 (bgg116)* and *LD2 (bgg212)*. **B**. Still frame images show impaired locomotion in *ubr-1* mutants swimming including altered tail movement (arrowheads). **C**. Schematic of MWT tracking for high-intensity locomotor swimming behavior. **D**. 30 sec MWT animal traces show individual *ubr-1* mutant with impaired locomotion. **E**. Quantitation of locomotor defects in *ubr-1* mutants and UBR-1 transgenic rescue. Shown are MWT plots of average locomotor speed (left) and expanded quantitation at 90 minutes (right) for indicated genotypes. **F**. AlphaFold prediction of UBR-1 structure. Inset shows RING domain (teal) and CRISR-engineered residues that are altered in *ubr-1 LD1* (dark blue) and *LD2* (green) mutants. Also shown are predicted coordination sites for Zn^2+^ ions (red circles), UBR box 1 (purple) and box 2 (orange). **G**. Quantitation indicates locomotor speed is reduced in *ubr-1 LD1* and *LD2* mutants. **For E and G**, MWT plots (solid lines, left) represent average speed of all recorded animals (4 animals/well, 4-5 wells per genotype per experiment, and 3-4 independent experiments) and shaded regions are SEM. Bars represent average of all wells for indicated time point, dots represent single wells tracked (4 animals/well), and error bars are SEM. Significance for plots (genotype annotations, left) tested with pairwise two-way ANOVA, and significance for bars with dots (right) tested using one-way ANOVA with Bonferroni’s post-hoc correction for multiple comparisons. **** p<0.0001. *** p<0.001

To quantitatively evaluate high-intensity locomotor behavior, we turned to multi-worm tracker (MWT). MWT is a computational behavioral tracking suite that allows unbiased, simultaneous monitoring of swimming speed for multiple *C. elegans* (**Fig 1C**) ^30^. MWT plots of swimming tracks for a single animal showed that *ubr-1 (tm5996)* mutants displayed reduced locomotion in liquid (**Fig 1D; Supplementary Fig 3**). Quantitation indicated that *ubr-1* mutants display robust, significant defects in locomotion compared to wild-type animals (**Fig 1E**). To confirm that UBR-1 regulates locomotor behavior, we used MosSCI to insert a single transgenic copy of UBR-1 into *ubr-1 (tm5996)* mutants. Transgenic UBR-1 expressed using the native promotor and 3’ UTR significantly rescued locomotor defects in mutants (**Fig 1E**).

Next, we tested whether *ubr-1* regulates locomotor behavior via RING ubiquitin ligase activity. To do so, we used CRISPR engineering to generate two *ubr-1* mutant alleles previously shown to abolish RING ubiquitin ligase activity ^31,32^. The first allele is *ubr-1 (bgg212)* carrying C1286A, H1288A and H1291A mutations, which we refer to as *ubr-1* ligase-dead 1 (*ubr-1 LD1*) (**Fig 1A**). The second is *ubr-1 (bgg116)*, carrying C1219S, which we refer to as *ubr-1 LD2* (**Fig 1A**). AlphaFold structural predictions illustrate how LD1 and LD2 both disrupt overlapping catalytic sites in the UBR-1 RING domain (**Fig 1F**). LD1 affects residues in two zinc-coordinating motifs in this location, while LD2 disrupts a single motif (**Fig 1F**). MWT was used to quantitatively evaluate swimming speed for these *ubr-1* LD1 and LD2 mutants. Both LD mutants displayed significantly decreased locomotor speed compared to wild-type animals, but also showed slightly milder locomotor defects than *ubr-1 (tm5996)* molecular null mutants (**Fig 1G**).

Thus, findings with three independently derived *ubr-1* loss-of-function alleles and transgenic rescue demonstrate that UBR-1 regulates high-intensity locomotor behavior. Mechanistically, our results suggest that UBR-1 utilizes RING ubiquitin ligase activity, and non-ligase mechanisms to shape high-intensity locomotor behavior.

### UBR-1 is expressed broadly in head neurons and functions in pre-motor interneurons to regulate locomotor behavior

To understand the cellular mechanism underpinning how UBR-1 regulates high-intensity locomotor behavior, we began by examining where UBR-1 is expressed using two approaches. First, we examined *ubr-1* expression in the *C. elegans* single-cell neural transcriptional atlas, CenGEN ^33^. Second, we used CRISPR to engineer a GFP fusion with endogenous UBR-1 (**Fig 2A**). Analysis of CenGEN data showed that *ubr-1* transcripts were detected in pre-motor interneurons, other interneurons, and sensory neurons which are all located in the head and tail of the animal (**Supplementary Table 1**). To visualize UBR-1 protein expression and localization, we used super-resolution microscopy on live, anesthetized *C. elegans*. GFP::UBR-1 expression was detected broadly across neurons in the head and tail (**Fig 2B&C; Supplementary Fig 4**). UBR-1 was localized to neuronal cell bodies, excluded from neuronal nuclei, and detected at lower levels in the axon bundle that forms the nerve ring (**Fig 2C**). 3D projections of super-resolution imaging were used to further visualize GFP::UBR-1 localization to the nerve ring and neuronal cell bodies in the head (**Supplementary Movies 5 and 6**).

**Fig 2:**
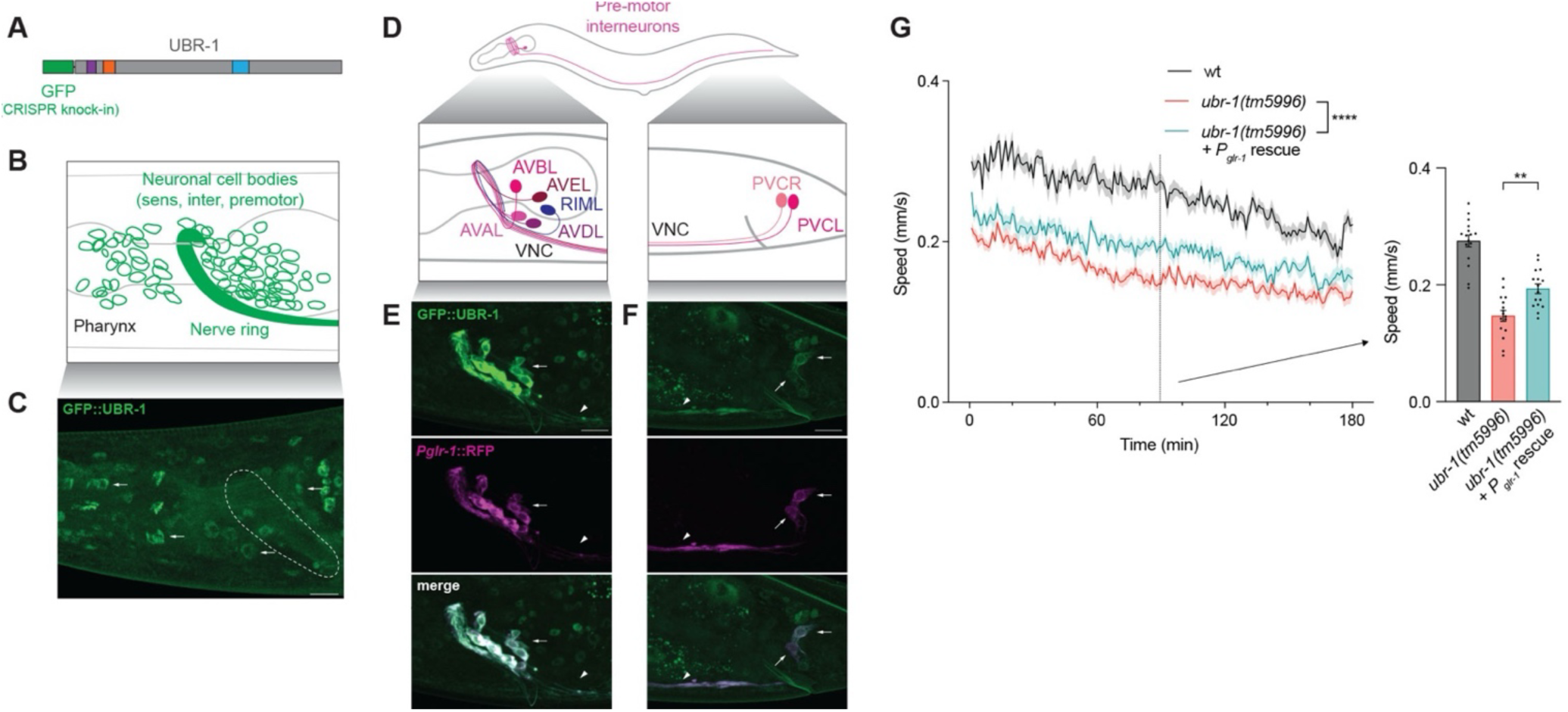
UBR-1 is expressed broadly in head neurons including pre-motor interneurons where it functions to regulate locomotor behavior. **A**. Schematic of GFP::UBR-1 generated by CRISPR engineering. **B**. Diagram of *C. elegans* head where cell bodies of sensory, interneurons and pre-motor interneurons as well as axonal nerve ring are located. **C**. Super-resolution images of adult *C. elegans* showing CRISPR engineered GFP::UBR-1 is expressed and localized to neuronal cell bodies (arrows) and nerve ring (dashed oval). **D**. Diagram shows pre-motor interneurons in head and tail of *C. elegans* with axons entering ventral nerve cord (VNC). Interneurons occur on both sides of head but only one side is depicted. **E and F.** Super-resolution images of GFP::UBR-1 (green) in cell bodies (arrows) and axons (arrowhead) of pre-motor interneurons in head (**E**) and tail (**F**). Pre-motor interneurons visualized using P*glr-1*::RFP (magenta). **G**. Quantitation of locomotor speed indicates UBR-1 expressed in pre-motor interneurons partially rescues locomotor defects in *ubr-1* mutants. **For G**, significance for data shown in plots (genotype annotations, left) tested with pairwise two-way ANOVA, and significance for bars with dots (right) tested using one-way ANOVA and Bonferroni’s post-hoc correction. ** p<0.01 Scale bar is 10 µm.

We further assessed GFP::UBR-1 expression and localization in neurons involved in locomotion using fluorescent reporters for pre-motor interneurons (*P_glr-1_*) and GABAergic motor neurons (*P_flp-13_*). GFP::UBR-1 was detected in the axons and cell bodies of pre-motor interneurons in the head and tail (**Fig 2D-F; Supplementary Figure 5**). In contrast, we did not detect GFP::UBR-1 in the cell bodies or axons of GABAergic motor neurons in ventral nerve cord or in the head (**Supplementary Fig 6**). We also did not detect GFP::UBR-1 in cell bodies of cholinergic motor neurons that are located directly adjacent to GABA neurons across the ventral cord (**Supplementary Fig 6**).

These findings on expression and localization suggested that UBR-1 could function in pre-motor interneurons, other interneurons or sensory neurons to regulate locomotor behavior. We tested this by performing single-copy transgenic rescue using the *glr-1* promoter (P*_glr-1_*), which expresses UBR-1 in pre-motor interneurons. This resulted in a partial, but significant, rescue of locomotor defects caused by *ubr-1 (tm5996)* (**Fig 2G**).

Collectively, our results support several conclusions. 1) Super-resolution microscopy detected endogenous CRISPR engineered GFP::UBR-1 in a variety of neurons in the head and tail. This is consistent with expression in sensory neurons, interneurons and pre-motor interneurons all of which contribute to upstream components of *C. elegans* sensorimotor circuitry. 2) UBR-1 localizes to cell bodies and axons of numerous neurons in the head including pre-motor interneurons. 3) Transgenic rescue results indicate that UBR-1 influences locomotor behavior, in part, by functioning cell-autonomously in pre-motor interneurons. These findings provide initial insight into the cellular mechanisms that UBR-1 utilizes to regulate locomotor behavior.

### CRISPR-based native proteomics indicates UBR-1 is a scaffold for a glutamate enzymatic network linked to human neurological abnormalities

Having examined neuronal mechanisms underpinning UBR-1 effects on locomotor behavior, we next delved into molecular mechanisms of UBR-1 function. To do so, we turned to unbiased affinity purification (AP) proteomics. Prior AP-proteomics studies in *C. elegans* that relied upon transgenic affinity tagged reagents have proven valuable for deciphering ubiquitin ligase signaling networks ^30,31,34^. Here, we increased the physiological relevance of AP-proteomics in *C. elegans* by deploying the CRISPR engineered endogenous GFP::UBR-1 strain that was used for localization and expression studies.

To ensure CRISPR engineered GFP::UBR-1 is functionally optimized for *C. elegans* proteomics, we evaluated GFP affinity tag placement on UBR-1 for effects on structure and function. AlphaFold structural predictions for *C. elegans* UBR-1 suggested that an N-terminal GFP tag was not folded near the UBR-1 catalytic site (**Supplementary Fig 7A**). In contrast, a C-terminal tag was predicted to fold close to the E2 and ubiquitin binding pockets, which could sterically hinder ubiquitin ligase activity (**Supplementary Fig 7B**). Consistent with these predictions, N-terminally tagged GFP::UBR-1 CRISPR engineered animals did not show locomotor defects (**Supplementary Fig 7C**), while C-terminally tagged UBR-1::GFP animals displayed similar locomotor defects to *ubr-1* mutants (**Supplementary Fig 7D**). Thus, both computational structural predictions and genetic results indicate that an N-terminal GFP tag does not interfere with UBR-1 folding and function, making it optimal for AP-proteomics (**Fig 3A**).

**Fig 3:**
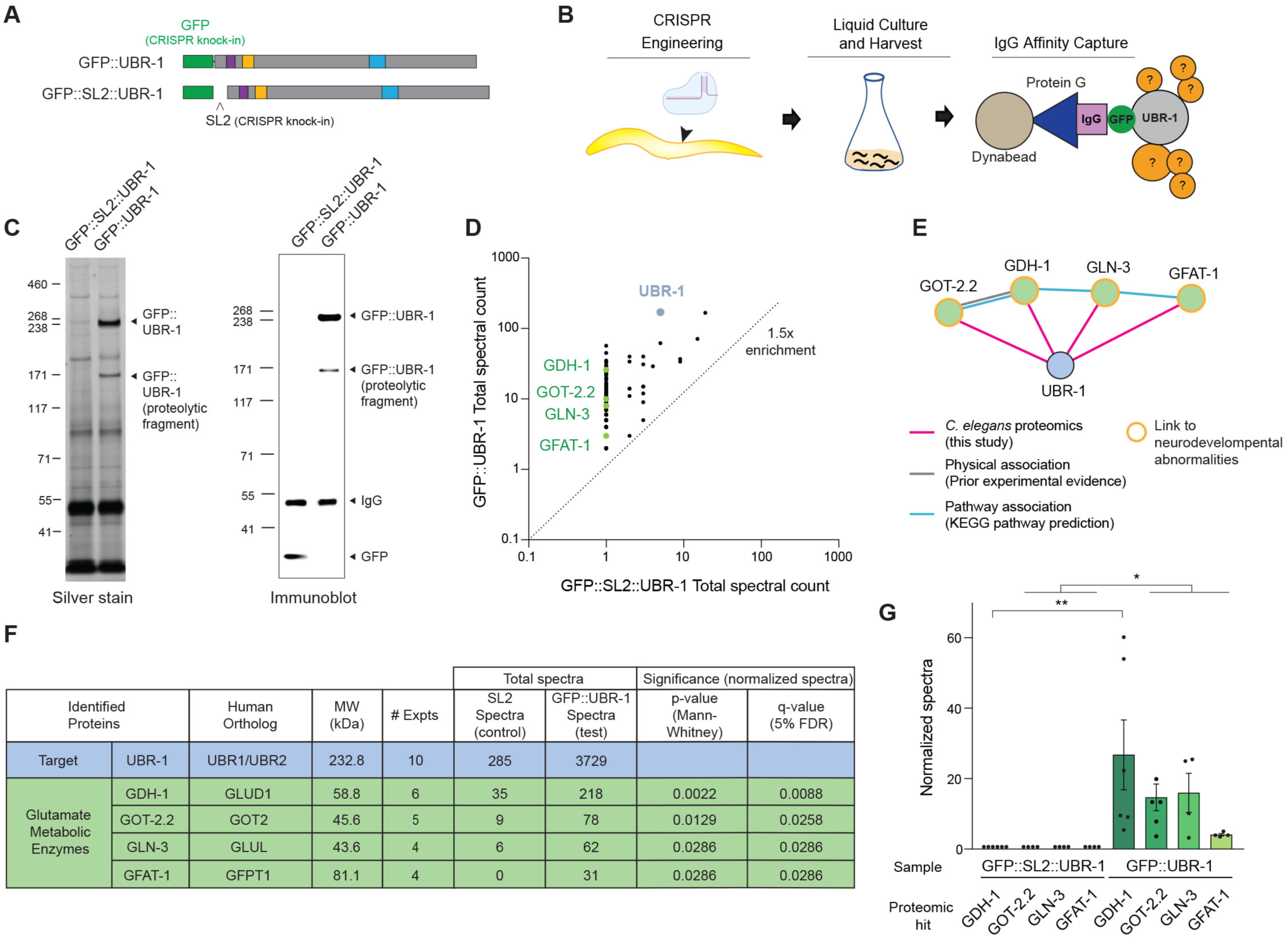
Large-scale CRISPR-based proteomics with UBR-1 identifies glutamate enzyme mini-network. **A**. Schematics depict GFP::UBR-1 and GFP::SL2::UBR-1 CRISPR engineered constructs used for AP-proteomics. **B**. Summary of CRISPR-based AP-proteomics workflow for *C. elegans*. **C**. Example of silver stain (left) and immunoblot (right) of anti-GFP purified samples used for proteomics. **D**. Example of single proteomics experiment with proteins identified for GFP::UBR-1 sample (test sample, y-axis) and GFP::SL2::UBR-1 sample (negative control, x-axis). Only protein hits with 1.5-fold or greater enrichment in GFP::UBR-1 sample (dotted line) were plotted. Highlighted proteins are UBR-1 affinity target (gray) and glutamate metabolic enzymes: GDH-1, GOT-2.2, GLN-3, AND GFAT-1 (green). **E**. Diagram of STRING computational network analysis for GOT-2.2, GDH-1, GLN-3, and GFAT-1. Pink lines highlight protein-protein interactions identified in this study between UBR-1 and glutamate enzyme mini-network. **F**. Summary of mass spectrometry results and statistics for 10 independent UBR-1 proteomics experiments from *C. elegans*. **G**. Quantitative analysis of glutamate enzyme enrichment in GFP::UBR-1 proteomics compared to GFP::SL2::UBR-1 (SL2) negative controls. **For F and G**, significance tested using Mann-Whitney and adjusted for multiple comparisons with 5% false-discovery rate (FDR). ** p<0.01, * p<0.05

Next, we proceeded with affinity purification of native GFP::UBR-1 from *C. elegans*. As a negative control, we CRISPR engineered an SL2 splice-leader with a stop codon between GFP and UBR-1 (GFP::SL2::UBR-1) (**Fig 3A**). This control strain independently expresses GFP and UBR-1 simultaneously via the native *ubr-1* promotor. AP-proteomics was performed as described previously (**Fig 3B**) ^35^. In brief, mixed stage *C. elegans* were grown in liquid culture, cleaned by sucrose flotation, and cryo-milled to generate submicron particles under liquid nitrogen temperatures. UBR-1 protein complexes were extracted and purified from whole worm lysates using anti-GFP antibodies coupled to Dynabeads. To ensure purification was optimized with GFP::UBR-1 sufficiently enriched in test samples compared to GFP::SL2::UBR-1 negative control samples, we performed both immunoblotting and silver staining on immunoprecipitates (**Fig 3C; Supplementary Fig 8**). We detected ∼232 kDa full-length GFP::UBR-1 and control GFP in immunoblots (**Fig 3C**). We also observed substantial enrichment of numerous unidentified proteins in SDS-PAGE silver stain gels of GFP::UBR-1 test samples compared to GFP::SL2::UBR-1 controls (**Fig 3C**). Thus, affinity purification from *C. elegans* lysates yielded robust enrichment of endogenous GFP::UBR-1 and UBR-1 protein complexes.

We performed large-scale proteomics analyses of affinity purified GFP::UBR-1 and UBR-1 protein complexes from *C. elegans* lysates in which ten independent, blinded experiments were run using three different detergent extraction conditions. Liquid chromatography tandem mass spectrometry (LC-MS/MS) or Trapped Ion Mobility Spectrometry – Time of Flight (TIMS-TOF) identified proteins in test and control samples. For analysis of mass spec data, proteins were considered hits if they were enriched at least 1.5-fold or higher in 2 or more experiments from GFP::UBR-1 samples compared to GFP::SL2::UBR-1 negative controls (**Fig 3D**). Across multiple proteomics experiments, we detected a mini-network of four glutamate metabolic enzymes that included: GDH-1, GFAT-1, GLN-3, and GOT-2.2 (**Fig 3D-F, Supplementary Table 2**). Using STRING and literature searches, we found no evidence that these enzymes were previously shown to form a complex with UBR-1 (**Fig 3E**). However, we did find published studies demonstrating that human orthologs of GDH-1 ^36,37^, GLN-3 ^38^, GOT-2.2 ^39^, and GFAT-1 ^40,41^ are all associated with human neurological deficits. The scale of our *C. elegans* proteomics analyses allowed us to further determine that all four glutamate enzymes were significantly enriched in GFP::UBR-1 samples compared to controls (**Fig 3F, G**). Thus, our analyses with large-scale, CRISPR-based proteomics suggest that UBR-1 is potentially a hub for a mini-network of glutamate metabolic enzymes that is associated with human neurodevelopmental deficits.

### UBR-1 interacts genetically with glutamate enzymes to regulate high-intensity locomotor behavior

With proteomics analyses suggesting UBR-1 could act mechanistically as a hub for glutamate enzymes, we wanted to test this model further using genetics. While the function of GDH-1, GFAT-1, GLN-3, and GOT-2.2 have not been studied in *C. elegans*, these glutamate metabolic enzymes are well characterized in other systems ^42^. Phylogenetic analysis with Alliance of Genome Resources (www.alliancegenome.org) and WormBase (wormbase.org) showed that GOT-2.2, GDH-1, GLN-3 and GFAT-1 are conserved from *C. elegans* to mammals. GOT-2.2 and GFAT-1 are predicted to catalyze glutamate synthesis from α-ketoglutarate and glutamine precursors, respectively (**Fig 4A; Supplementary Fig 9**). In contrast, GLN-3 and GDH-1 are predicted to facilitate conversion of glutamate to glutamine or α-ketoglutarate. Here, we generated molecular null alleles of glutamate metabolic enzymes to evaluate their effects on high-intensity locomotor behavior and examine functional interactions with UBR-1.

**Fig 4:**
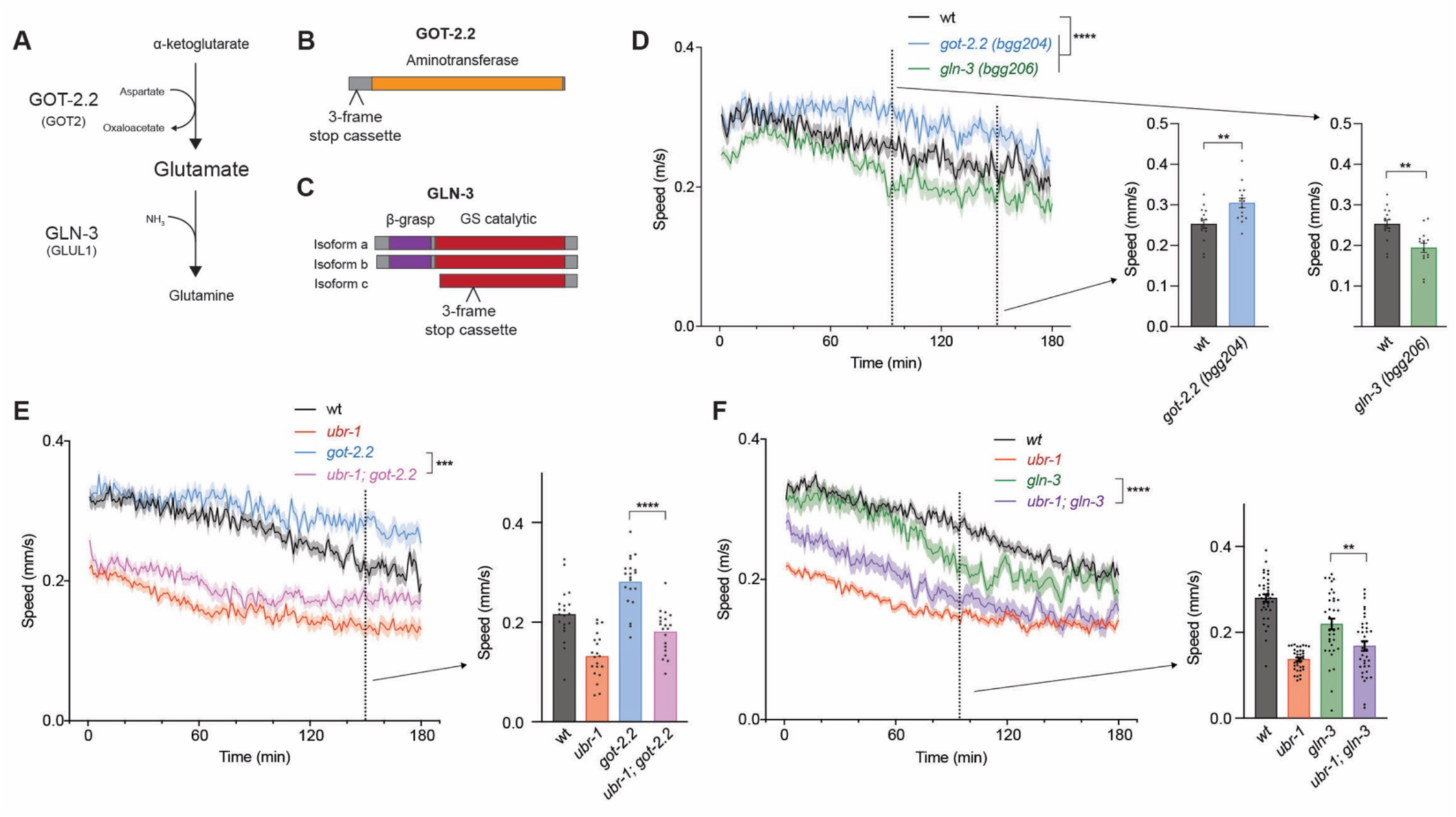
GOT-2.2/UBR-1/GLN-3 axis regulates locomotor activity. **A**) Schematic showing opposing effects of GOT-2.2 and GLN-3 on glutamate metabolism. **B and C**. Protein diagrams of GOT-2.2 and GLN-3 isoforms with CRISPR engineered three-frame STOP cassettes. **D**. Quantitation of locomotor swimming speed showing sustained locomotion in *got-2.2(bgg204)* mutants and early fatigue in *gln-3(bgg206)* mutants in MWT plots (left). Expanded quantitation is shown for *got-2.2* at 150 mins (middle) and *gln-3* at 95 mins (right). **E**. Quantitation shows *ubr-1; got-2.2* double mutants have similar decrease in speed as *ubr-1* mutants, and significant decrease compared to *got-2.2* single mutants. **F**. Quantitation shows *ubr-1; gln-3* double mutants have similar decrease in speed to *ubr-1* single mutants. **For D-F**, MWT plots (solid lines, left) represent average speed of all recorded animals (4 animals/well, 5 wells per genotype per experiment, and 3-4 independent experiments) and shaded regions are SEM. Bars (right) represent average of all wells for indicated time point, dots represent single wells tracked (4 animals/well), and error bars are SEM. Significance for plots (genotype annotations, left) tested with pairwise two-way ANOVA, and significance for bars with dots (right) tested using one-way ANOVA with Bonferroni’s correction for multiple comparisons. **** p<0.0001. *** p<0.001, ** p<0.01

Because GLN-3 and GOT-2.2 have opposing effects on glutamate metabolism (**Fig 4A**), we hypothesized these enzymes could have differential effects on *C. elegans* locomotor behavior. To test this, we generated molecular null alleles for glutamate metabolic enzymes using CRISPR to knock in three-frame stop cassettes ^43^. We failed to isolate null alleles for *gdh-1* and *gfat-1*, which is likely due to lethality. However, we did obtain multiple molecular null alleles for both *got-2.2* (*bgg204, bgg205*) and *gln-3* (*bgg206, bgg207*) (**Fig 4B & C**). Consistent with differing roles in glutamate metabolism, we observed opposing effects on swimming speed for *got-2.2* and *gln-3* mutants (**Fig 4D**). Both *got-2.2* mutant allele isolates resulted in significant sustained swimming with failure to fatigue as rapidly as wild type (**Fig 4D; Supplementary Fig 10A**). In contrast, both *gln-3* mutant isolates displayed the opposite phenotype, early fatigue (**Fig 4D**;

**Supplementary Fig 10B**). These results indicate that both GOT-2.2 and GLN-3 affect high-intensity locomotor behavior, similar to UBR-1.

To evaluate genetic interactions between glutamate enzymes and *ubr-1*, we used CRISPR to directly engineer the same molecular nulls for *got-2.2 (bgg208)* and *gln-3 (bgg210)* into *ubr-1 (tm5996)* deletion mutants. While these are the same three-frame stop cassette insertions as *gln-3 (bgg206)* and *got-2.2 (bgg204)* single mutants, they are designated as separate alleles because they were engineered independently. We found that *ubr-1; got-2.2* double mutants displayed a similar decrease in swimming speed as *ubr-1* single mutants (**Fig 4E**). The decrease in speed in double mutants was significant and the opposite phenotype to what occurs in *got-2.2* single mutants (**Fig 4E**). Our findings that *ubr-1* and *got-2.2* single mutants display opposing locomotor defects with predominantly reduced swimming speed (*i.e. ubr-1* like defects) in double mutants is consistent with GOT-2.2 being an upstream inhibitor of UBR-1. With regard to *gln-3*, single mutants displayed reduced swimming speed similar to *ubr-1* single mutants (**Fig 4F**). For *ubr-1; gln-3* double mutants, we found similar locomotor defects to what was observed in *ubr-1* single mutants (**Fig 4F**). Because we are using molecular null alleles for *ubr-1* and *gln-3*, these results indicate that UBR-1 and GLN-3 function in the same genetic pathway to regulate locomotor behavior. Taken together, our genetic interaction results and proteomic findings indicate that UBR-1 is a hub in a GOT-2.2/UBR-1/GLN-3 axis that is required for high-intensity locomotion in *C. elegans*.

### GOT-2.2/UBR-1/GLN-3 axis regulates glutamate homeostasis to influence locomotor behavior

Next, we wanted to further evaluate the cellular mechanism for how UBR-1 functionally integrates with glutamate metabolic enzymes in the GOT-2.2/UBR-1/GLN-3 axis to influence locomotor behavior. Our results indicate UBR-1 is expressed in interneurons and sensory neurons, and functions cell-autonomously in interneurons (**Fig 2; Supplementary Table 1**). Both interneurons and sensory neurons rely upon glutamatergic signaling and metabolism, and function upstream of motor neurons in *C. elegans* sensorimotor circuitry (**Fig 5A**) ^44,45^. During locomotion, upstream glutamate signaling converges on excitatory motor neurons, which signal via acetylcholine (ACh) to stimulate muscles to contract. GABA motor neurons inhibit muscles to generate relaxation. Opposing ACh excitation and inhibitory GABA motor neuron function is required to produce high-intensity locomotor behavior. If the GOT-2.2/UBR-1/GLN-3 axis is required to maintain glutamate homeostasis, we would expect these players to impact downstream ACh motor neuron function thereby influencing locomotor behavior (**Fig 5A**).

**Fig 5:**
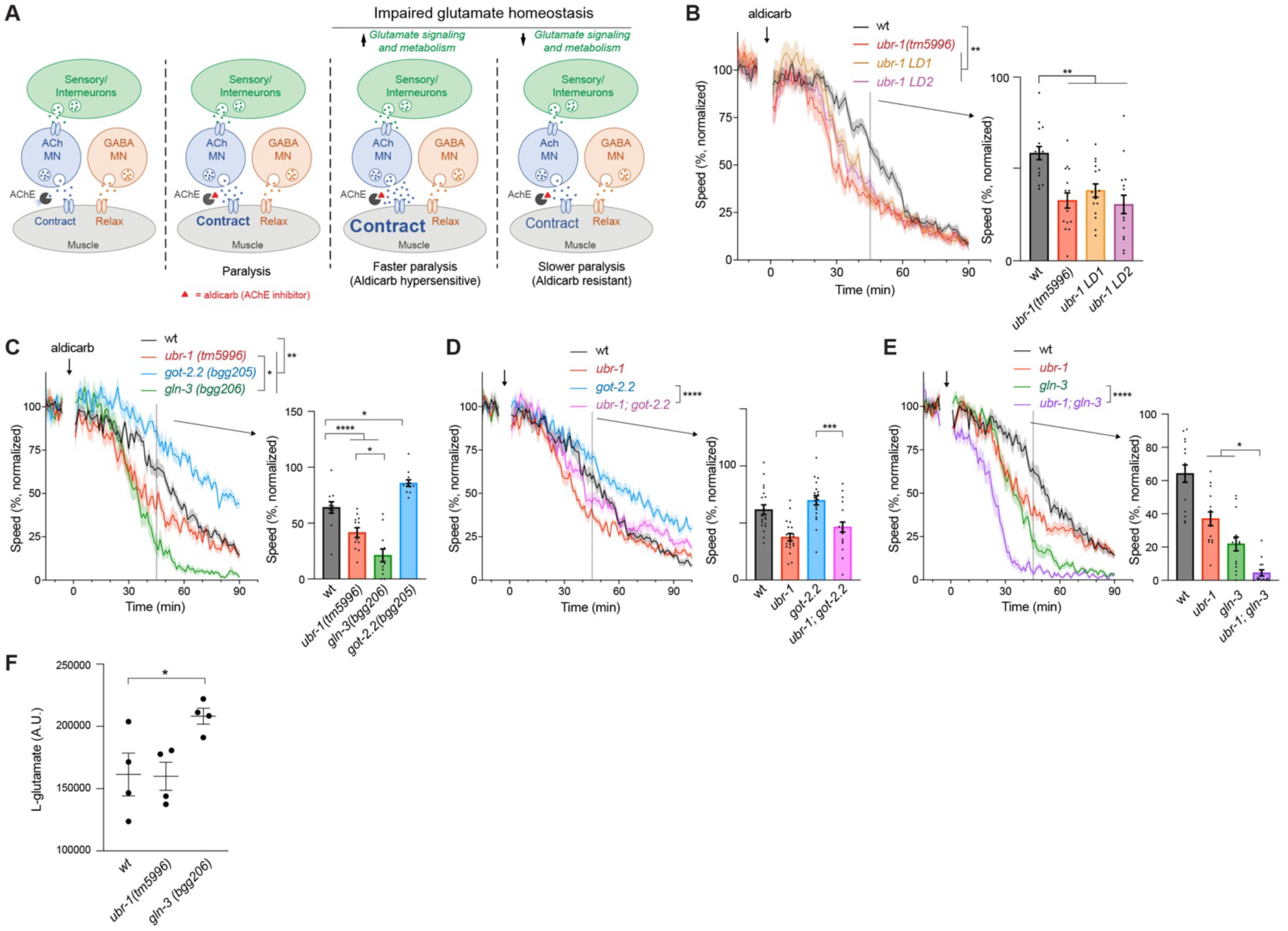
GOT-2.2/UBR-1/GLN-3 axis regulates glutamate homeostasis to affect cholinergic motor neurons and locomotor behavior. **A**. Simplified *C. elegans* sensorimotor circuit showing how aldicarb acetylcholine esterase (AChE) inhibitor alters ACh motor neuron function. Right panels depict how altering glutamate homeostasis influences ACh motor neuron function and aldicarb sensitivity. **B**. Quantitative MWT results show *ubr-1* mutants are hypersensitive to aldicarb treatment (arrow). Shown are normalized MWT plots of average locomotor speed (left) and expanded quantitation at 45 mins (right). **C**. Quantitative results show *got-2.2* mutants are aldicarb resistant, and both *gln-3* and *ubr-1* mutants display aldicarb hypersensitivity. **D**. Quantitation shows aldicarb resistance is suppressed in *ubr-1; got-2.2* double mutants compared to *got-2.2* single mutants. **E**. Quantitation shows aldicarb hypersensitivity is enhanced in *ubr-1; gln-3* double mutants compared to *gln-3* single mutants. **F**. Quantitative LC-MS results show increased whole-animal glutamate levels in *gln-3* mutants. **For B-E**, MWT plots (solid lines, left) represent average speed of all recorded animals (4 animals/well, 5 wells per genotype per experiment, and 3-4 independent experiments) and shaded regions are SEM. MWT plots are normalized to average speed over 10 min baseline prior to aldicarb treatment. Bars (right) represent average of all wells for indicated time point, dots represent single wells tracked (4 animals/well), and error bars are SEM. Significance for plots (genotype annotations, left) tested with pairwise two-way ANOVA, and significance for bars with dots (right) tested using one-way ANOVA and Bonferroni’s post-hoc correction for multiple comparisons. **For F**, lines are mean, dots are single LC-MS result, and error bars are SEM. Significance determined using Student’s *t* test with Bonferonni correction **** p<0.0001. *** p<0.001, ** p<0.01, * p<0.05

To begin testing this hypothesis, we pharmacologically targeted cholinergic motor neurons using aldicarb, an acetylcholinesterase inhibitor (**Fig 5A**). Aldicarb blocks cholinesterase activity and causes a buildup of ACh, which over time leads to excess activation of cholinergic synapses, muscle hypercontraction, and eventual paralysis (**Fig 5A**). Aldicarb can be used to assess whether mutants of interest affect cholinergic or GABA motor neuron function by evaluating how long it takes for aldicarb-induced paralysis to occur.

We previously established methods for using MWT to monitor the effects of aldicarb in high-intensity locomotor swimming assays ^30^. If *ubr-1*, *gln-3* or *got-2.2* mutants have altered glutamate homeostasis in interneurons and/or sensory neurons, we would expect homeostatic effects on downstream cholinergic motor neuron function (**Fig 5A**). As a result, altered sensitivity to aldicarb would occur depending upon whether glutamate homeostasis was increased or decreased. We found that all three *ubr-1* mutants (*tm5996*, *LD1*, *LD2*) displayed faster reductions in swimming speed when treated with aldicarb compared to wild-type animals (**Fig 5B**). Thus, loss of UBR-1 function results in aldicarb hypersensitivity, which is consistent with increased glutamate homeostasis.

To more directly evaluate how changes in glutamate homeostasis affect cholinergic motor neuron function, we tested aldicarb sensitivity of *got-2.2* and *gln-3* mutants. We found that *got-2.2* mutants are significantly resistant to aldicarb-induced reduction in swimming speed compared to wild-type animals (**Fig 5C**). In contrast, *gln-3* mutants display aldicarb hypersensitivity (**Fig 5C**). The opposing effects of GLN-3 and GOT-2.2 on glutamate homeostasis evaluated via aldicarb is in line with their opposing effects on locomotor behavior in the absence of aldicarb.

Next, we assessed aldicarb responses of *ubr-1 (tm5996)* double mutants with *got-2.2* or *gln-*3. For *ubr-1; got-2.2* double mutants, we observed a significant suppression of aldicarb resistance seen in *got-2.2* single mutants (**Fig 5D**). This is consistent, with *ubr-1* suppressing *got-2.2* mutant phenotypes in locomotor studies without aldicarb (**Fig 4E**. In contrast, aldicarb hypersensitivity was prominently and significantly enhanced in *ubr-1; gln-3* double mutants compared to *gln-3* or *ubr-1* single mutants (**Fig 5E**). These results suggest that UBR-1 and GLN-3 function coordinately to regulate glutamate homeostasis.

To further evaluate the model that GLN-3 and UBR-1 affect glutamate homeostasis, we turned to liquid chromatography mass spectrometry (LC-MS). Using LC-MS, we quantitatively evaluated whole organism levels of glutamate. We observed a significant increase in glutamate levels in *gln-3* mutants compared to wild type (**Fig 5F**). This finding is consistent with GLN-3 mediating the metabolic transition of glutamate to glutamine (**Fig 4A**). In prior studies ^27^, *ubr-1* mutants were found to have increased glutamate levels. However, our LC-MS metabolomics analyses for *ubr-1 (tm5996)* mutants did not yield a significant increase in glutamate levels (**Fig 5F**). Importantly, more substantial glutamate alterations in *gln-3* mutants are consistent with more severe aldicarb hypersensitivity in *gln-3* mutants compared to *ubr-1* mutants (**Fig 5C**).

Taken as a whole, our findings support several conclusions. 1) Our results from single mutants and multi-gene interaction studies with aldicarb assays provide further support for UBR-1, GLN-3 and GOT-2.2 functioning in a genetic axis. 2) UBR-1 and GLN-3 have a positive genetic interaction that affects glutamate homeostasis, which is revealed by pharmacological pressure on cholinergic motor neurons. 3) Finally, all our findings are consistent with the GOT-2.2/UBR-1/GLN-3 axis affecting glutamate homeostasis to influence high-intensity locomotor behavior.

### UBR-1 is expressed in gonad and functions with GLN-3 to regulate animal viability

Because JBS patients have developmental delay, we expanded our studies to evaluate whether UBR-1 affects organismal development. To inform our choice of phenotypic readouts, we initially used super-resolution microscopy to examine where CRISPR engineered GFP::UBR-1 is expressed outside of the nervous system. Interestingly, we observed GFP::UBR-1 expression in germ cells, oocytes, and embryos (**Fig 6A-C, Supplementary Fig 11)**. Similar to neurons, UBR-1 was excluded from the nucleus. This expression pattern suggests UBR-1 could play a role throughout early animal development.

**Fig 6:**
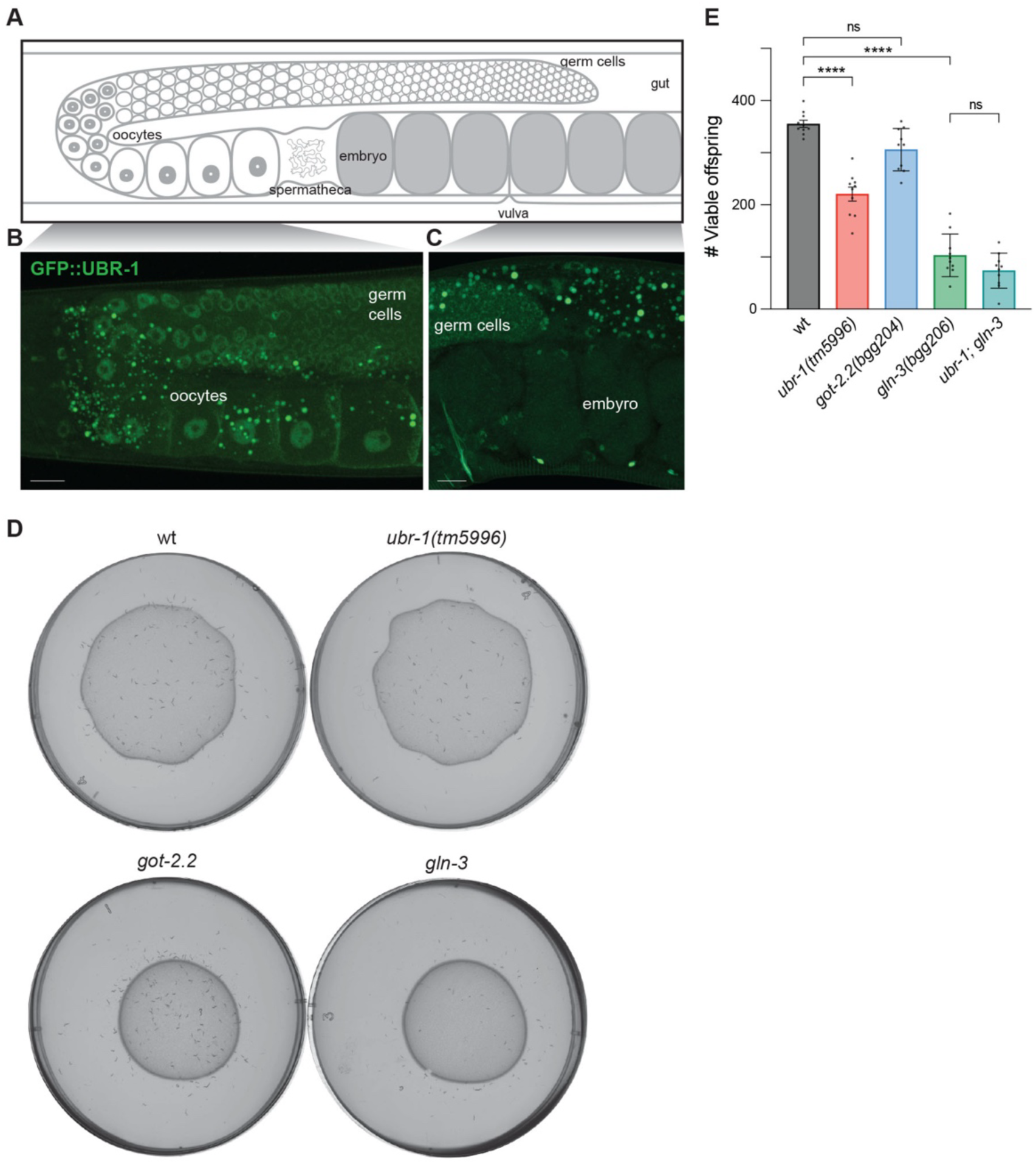
UBR-1 is expressed in gonad and functions with GLN-3 to regulate developmental viability. **A**. Diagram of *C. elegans* adult gonad with germ cells, oocytes and embryos. **B and C**. Super-resolution images showing CRISPR engineered GFP::UBR-1 is expressed in germ cells and oocytes (**B**) as well as germ cells and embryos (**C**). **D**. Representative plates showing offspring laid by a single adult animal after 96 hrs for indicated genotypes. **E**. Quantitation shows reduced total viable offspring for *ubr-1* and *gln-3* mutants compared to wt. Viability is not further reduced in *ubr-1(tm5996); gln-3* double mutants compared to *gln-3* single mutants. Dots represent total viable offspring laid over 96 hrs from a single adult. Significance was tested with one-way ANOVA and Bonferroni’s post-hoc correction for multiple comparisons. **** p<0.0001, ns = not significant Scale bar is 10 µm

We next performed viability assays on *ubr-1 (tm5996)* mutants. We observed a significant reduction in total viable offspring laid by *ubr-1* mutants compared to wild-type hermaphrodites (**Fig 6D&E**). To examine whether changes in glutamate metabolism contribute to viability, we evaluated *gln-3* and *got-2.2* mutants. We found that *gln-3* mutants showed prominent reductions in animal viability compared to wild-type animals (**Fig 6D&E**). In contrast, we observed no effects on viability for *got-2.2* mutants (**Fig 6E**).

With our results suggesting both UBR-1 and GLN-3 play a role in animal viability, we next examined *ubr-1; gln-3* double mutants. Interestingly, *ubr-1; gln-3* double mutants did not show increased viability defects compared to *gln-3* single mutants (**Fig 6E**). Because we are using molecular null alleles, our observations indicate that UBR-1 and GLN-3 function in the same pathway to regulate animal viability. Thus, results from developmental viability studies further reinforce genetic conclusions from locomotor studies. Overall, our findings support the model that UBR-1 affects glutamate homeostasis to influence both locomotor behavior and developmental viability (**Fig 7**).

**Fig 7:**
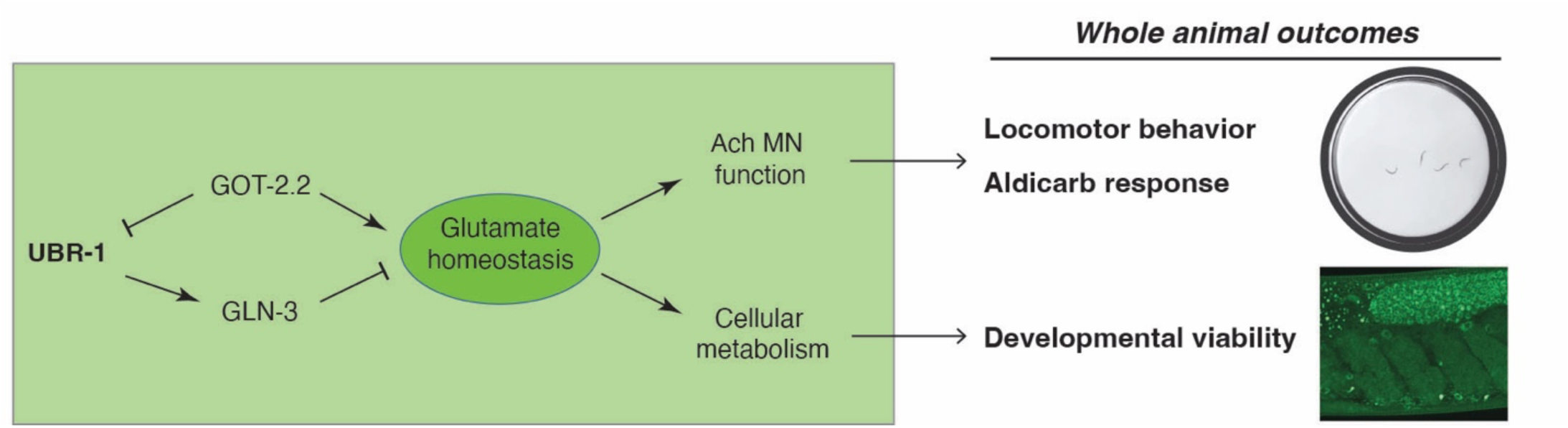
GOT-2.2/UBR-1/GLN-3 axis regulates glutamate homeostasis to affect high-intensity locomotor behavior and developmental viability. Summary shows that UBR-1 functions as an enzyme hub in the GOT-2.2/UBR-1/GLN-3 axis that is required to maintain glutamate homeostasis. UBR-1 axis regulation of glutamate homeostasis affects ACh motor neuron function and cellular metabolism leading to outcomes on locomotor behavior, sensitivity to aldicarb perturbation of ACh motor neuron function, and developmental viability.

## Discussion

Our investigation has shown that UBR-1 regulates high-intensity locomotor behavior and organismal viability. Both these phenotypes share similarities with global developmental delay and failure to reach motor development milestones reported for JBS patients. Consistent with our phenotypic observations in *C. elegans*, super-resolution imaging indicates that UBR-1 is expressed and localized to both axons and soma of interneurons that are influenced by glutamate metabolism and signaling, as well as in germ cells, oocytes, and developing embryos. We combined CRISPR-based proteomics and genetics to delve into the molecular genetic mechanisms by which UBR-1 regulates locomotor behavior and viability. Our results indicate that UBR-1 acts as an enzyme hub for a mini-network of glutamate metabolic enzymes including GLN-3, GOT-2.2, GFAT-1 and GDH-1. Genetic and pharmacological results indicate that functional integration of the GOT-2.2/UBR-1/GLN-3 axis is required for balanced glutamate homeostasis that mediates high-intensity locomotion and organismal viability (**Fig 7**).

Prior yeast and mammalian cell-based studies have revealed much about the structure and biochemistry of UBR1 and its mechanisms of substrate recognition ^11,32,46,47^. Work in these cellular models has also shown UBR1 regulates lipid homeostasis involved in hepatic and muscle steatosis ^48^. Several of our findings now provide key support in an organismal model that UBR-1 functions as an enzyme hub that regulates glutamate homeostasis to affect locomotor behavior. 1) Our expression studies and genetic rescue experiments demonstrate native UBR-1 is expressed and functions in head neurons, including pre-motor interneurons which utilize glutamate signaling (**Fig 2**). 2) Proteomic and genetic results indicate that UBR-1 is a scaffold for multiple glutamate metabolic enzymes and functionally integrates with GLN-3 and GOT-2.2 (**Fig 3&4**). 3) Our behavioral pharmacology findings indicate that UBR-1, GOT-2.2, and GLN-3 are each necessary for efficient locomotor activity (**Fig 5**). 4) Finally, prior orthogonal studies found that UBR-1 can regulate reverse locomotor behavior and ivermectin resistance via a fifth enzyme that affects glutamate metabolism, GOT-1 ^49,50^. Collectively, these findings indicate that UBR-1 has a multi-faceted physical and genetic relationship with numerous glutamate metabolic enzymes in *C. elegans*. Thus, we conclude that UBR-1 is an enzyme hub for a metabolic mini-network that is required to maintain glutamate homeostasis **(Fig 7**).

Importantly, our genetic results do not support the model that UBR-1 polyubiquitinates and inhibits GOT-2.2 or GLN-3 through an enzyme-substrate relationship. Consistent with this, we did not identify an N-degron consensus sequence in either GLN-3 or GOT-2.2 protein sequences (**Supplementary Fig 12**). Our findings with molecular null alleles across a range of phenotypic readouts indicate that GOT-2.2 is likely to function as an upstream inhibitor of UBR-1 (**Fig 4 & 5**). In contrast, GLN-3 functions in the same pathway with UBR-1 and is potentially downstream and positively regulated by UBR-1. However, our observations with CRISPR-engineered *ubr-1 LD* mutants suggest that UBR-1 ubiquitin ligase activity is also required for locomotor behavior (**Fig 1**). Thus, our results suggest that UBR-1 functions as an enzyme hub and ubiquitin ligase that integrates the function of glutamate metabolic enzymes to influence locomotion. Importantly, other ubiquitin ligase signaling hubs associated with childhood neurodevelopmental disorders, such as RPM-1 (MYCBP2) and EEL-1 (HUWE1), also utilize both ubiquitin ligase activity and ligase-independent mechanisms to shape neuron development and function ^51,52^.

Our discovery that UBR-1 acts as a hub in the GOT-2.2/UBR-1/GLN-3 axis to regulate glutamate homeostasis provides important insight into the molecular and genetic mechanisms regulating locomotor behavior and developmental viability. Interestingly, all four glutamate metabolic enzymes detected in UBR-1 proteomics, including GOT-2.2 and GLN-3, have mammalian orthologs with documented human genetic variants associated with neurological phenotypes. 1) GOT-2.2 corresponds to human GOT2 mitochondrial aspartate transaminase. Bi-allelic loss of *GOT2* results in epilepsy and intellectual disability, and was recently termed developmental and epileptic encephalopathy-82 ^39^. 2) GLN-3 is orthologous to glutamine synthetase GLUL (also called GS), which is present in mammalian astrocytes and known to be linked to mesial temporal-lobe epilepsy ^38^. 3) GFAT-1 is the ortholog of human GFPT1, a key enzyme in the hexosamine pathway that is associated with myasthenic syndrome ^40,41^. Myasthenic syndrome features both neuromuscular weakness and leukoencephalopathy. 4) Finally, GDH-1 corresponds to human GLUD1, which is a glutamate dehydrogenase. Human genetic variants in *GLUD1* cause hyperinsulinism–hyperammonemia syndrome, which features neurological symptoms such as seizure and intellectual disability ^36,37^. Thus, UBR-1 modulation of this glutamate mini-network, glutamate enzyme links to neurological deficits, and UBR-1 involvement in JBS suggests this network could represent an evolutionarily conserved mechanism for safeguarding excitatory glutamatergic signaling.

Prior studies have examined *Ubr1* knockout mice and *Ubr1; Ubr2* double knockout mice primarily identifying pancreatic defects, reduced body weight and embryonic lethality ^2,12,13^. With regard to the nervous system, earlier findings indicate that neurogenesis is impaired in *Ubr-1; Ubr2* double knockout mice, with potential links to notch signaling ^13^. Our results suggest that more in depth studies on glutamate homeostasis, transmission, signaling and metabolism in *Ubr1* and *Ubr-1; Ubr2* knockout mice could be valuable. Importantly, findings across model systems indicate that excess glutamate signaling causes neuronal excitotoxicity ^17^. Elevated glutamate is also toxic in non-neuronal cell types that express glutamate receptors, such as pancreatic β-cells ^53^. Thus, GOT-2.2/UBR-1/GLN-1 axis effects on glutamate homeostasis identified here might have broader implications for the development and health of the nervous system as well as non-nervous tissue. As a result, it is possible UBR1 effects on glutamate homeostasis could contribute to both neurodevelopmental abnormalities and pancreatic insufficiency in patients with JBS.

## Methods

### Genetics and strains

The *C. elegans* N2 isolate was used to generate all strains used, and animals were maintained using standard procedures. The following mutant and CRISPR alleles were used: *ubr-1 (tm5996)* I, GFP::UBR-1 CRISPR (*bgg110*) I, UBR-1::GFP CRISPR (*bgg112*) I, GFP::SL2::UBR-1 CRISPR (*bgg140*) I*, ubr-1* LD1 CRISPR (*bgg212*) I*, ubr-1* LD2 CRISPR (*bgg116*) I, *gln-3* 3-frame stop CRISPR (*bgg206*, *bgg207*) IV, *got-2.2* 3-frame stop CRISPR (*bgg204*, *bgg205*) X. Double mutants were generated by CRISPR editing into *ubr-1 (tm5996)* to generate *ubr-1 (tm5996)*; *got-2.2* (*bgg208*) and *ubr-1 (tm5996); gln-3* (*bgg210*). The following transgenic strains were used: *odIs6* [*Pglr-1*::mRFP], *bggIs6* [*Pflp-13*::mCherry]. The following MosSCI strains were used: *bggSi56*[P_ubr-1_::UBR-1::*ubr-1* 3’UTR, *unc-119(+)*] II, *bggSi57*[P_glr-1_::UBR-1::*let-858* 3’UTR, *unc-119(+)*] II. MosSCI strains were inserted into *ttTi5605*. MosSCI strains were mated with *ubr-1 (tm5996)* to generate rescue strains: *ubr-1 (tm5996)*; *bggSi56*[P_ubr-1_::UBR-1], and *ubr-1 (tm5996)*; *bggSi57*[P_glr-1_::UBR-1]. All strains were grown at 20°C for at least 2 generations prior to executing behavioral experiments. Mutants, CRISPR alleles and transgenes used for specific experiments are listed in **Supplementary Table 3**. Primers used for strain genotyping are listed in **Supplementary Table 4**. Plasmids used to generate transgenic strains are listed in **Supplementary Table 5**. CRISPR reagents are listed in **Supplementary Table 6**. Microinjection conditions used to generate strains are shown in **Supplementary Table 7**.

### CRISPR/Cas9 engineering

CRISPR alleles were engineered via direct injection of Cas9 ribonucleoprotein complexes with *dpy-10* co-CRISPR. Complexes are formed by mixing recombinant Cas9 protein, tracrRNA (IDT), and crRNA (IDT) at 37 °C for 15 min, followed by addition of ssODN repair template (IDT). CRISPR-Cas9 ribonuclear complexes were injected into N2 animals, individual edited alleles were isolated by PCR-based genotyping, and confirmed by sequencing. STOP alleles were designed using a three-frame stop cassette ^54^, which was inserted into the earliest exon that affects all isoforms.

Repair template for GFP::SL2::UBR-1 was amplified from genomic DNA from *bgg110* (GFP::UBR-1 CRISPR) using Q5 High-Fidelity DNA Polymerase (NEB) system. Fragments were subcloned into pBG-98 using HiFI DNA assembly (NEB). Repair template was amplified, assembled, and injected as described previously ^55^. All CRISPR targeting sequences and repair templates are listed in **Supplementary Table 6**.

### High-intensity locomotor behavior and liquid aldicarb assays

We often use the term high-intensity locomotor behavior instead of swimming to make a clear distinction about the level of activity in our liquid-based behavioral locomotor experiments compared to locomotor studies on solid media where animals display less vigorous locomotion. Multi-worm tracker (MWT) was used to quantitatively monitor high-intensity locomotor swimming behavior as described previously ^30,56^. In brief, worms were synchronized by picking L4 animals 20-24 hrs prior to MWT assays. Young adult animals were picked into the lid of a 96-well plate. Lid wells contained 20 μl of assay buffer (M9 + 0.01% tween-20). MWT was used to record animal swimming speed for up to 3 hrs. MWT aldicarb assays were performed as described previously ^57^. For all experiments and genotypes, 10 min baseline was initially recorded and 20 μl of 100 µM aldicarb was added to wells to a final concentration of 50 µM. Swimming speed was recorded for up to 3 hrs. Analysis of locomotor speed was performed using a custom script, with speed defined as the movement speed of the animal centroid in mm/s. Data were analyzed and graphed using Prism (GraphPad)

### Video capture of C. elegans behavior

Movies of *C. elegans* on solid media plates and in liquid culture were captured on a Zeiss Stemi 305 stereoscope with Axiocam 208 color camera attachment. For liquid culture, plates were set up as noted above for high-intensity locomotor assays with 2 adjacent wells using 20 µl assay buffer and 4 animals per well. Liquid locomotion was captured for 5 minutes. For solid media, adult animals were transferred to fresh plates and allowed to crawl on plates for 2 minutes, and video was then captured for 2 minutes. Movies were annotated and clipped in Adobe Premiere Pro.

### AlphaFold predictions

Structures of UBR-1 with and without CRISPR engineered GFP were predicted using the LocalColabFold version of ColabFold 1.5.0 using pdb templates with “—templates” and relaxed using “—amber” ^58^ or AlphaFold3 ^59^. Five models were predicted for each protein and the model with the highest predicted local distance difference test (plDDT) was presented for each protein. Predicted structures were adjusted and colorized using Open-source PyMOL (PyMOL Molecular Graphics System, Version 2.5.0, Schrödinger, LLC).

### Super-resolution imaging of UBR-1 expression and localization

GFP::UBR-1 (*bgg110*) CRISPR engineered animals were used to evaluate UBR-1 expression and localization. For cell-specific studies, GFP::UBR-1 (*bgg110*) males were mated with hermaphrodites for *odIs6* [P_glr-1_::mRFP] or *bggIs6* [P_flp-13_::mCherry] which label premotor interneurons and GABA motor neurons, respectively. Images were then collected from *bgg110*; *odIs6* or *bgg110; bggIs6* animals. Super-resolution images were captured using a Zeiss LSM 980 with Airyscan 2. For imaging, animals were mounted onto slides with a thin layer of 3% agarose and immobilized using 5-10 mM levamisole. Images were acquired using 40x objective for all images.

3D movies of super-resolution imaging were generated using ZEN 3D Toolkit. Movies represent rotating z-stack generated from 80-120 super-resolution slices from wt or GFP::UBR-1 (*bgg110*) animals at slice thickness of 0.19 µm.

### Developmental viability assay

To evaluate organismal viability during development, single parent animals were transferred at L4 larval stage onto 6-cm plates with OP50 *E. coli*. Every 24 hours, parent was transferred to fresh plate. Progeny were considered viable if they reached L4 larva stage or older.

### C. elegans CRISPR-based native proteomics

*C. elegans* anti-GFP affinity purification from CRISPR engineered strains: CRISPR-based native proteomics was performed based on modification of previous AP-proteomics methods for *C. elegans* ^35^. Mixed-stage GFP::UBR-1 (*bgg110*) or GFP::SL2::UBR-1 (*bgg140*) animals were cultured for 2–3 days in flasks that contained S complete media, cholesterol and HB101 *E.coli*. Liquid cultures were mixed and aeriated on an orbital shaker (185 RPM). Worms were harvested by low-speed centrifugation (350 ⨉ g), separated on 30% sucrose flotation, and washed three times with 0.1M NaCl. Packed worms were frozen in pellets using liquid N2. Grindates were generated by cryomilling frozen worm pellets into submicron particles under liquid nitrogen cooling (Retsch) with EDTA-free protease inhibitor tablets (Roche). *C. elegans* grindates were lysed and extracted using four times volume lysis buffer (50mM Tris HCl (pH 7.5), 150mM NaCl, 1.5mM MgCl2, 10% glycerol, detergent [0.1% NP-40, 0.1% CHAPS, or 0.1% Triton X-100], 1mM DTT, 1mM PMSF, 1x HALT Protease Inhibitor Cocktail (Thermo Scientific), 1mM sodium orthovanadate, 5mM sodium flouride, 1mM sodium molybdate). Whole worm lysates were stirred for 2 min, rotated for 5 min at 4 °C, and cleared by high-speed centrifugation at 20,000×g (10 min). Affinity purification was performed using lysate volume equivalent to 20 mg total protein. Affinity purification targets were captured with 500 μL of Dynabeads (M280 anti-rabbit IgG, Invitrogen) under rotation for 4h at 4 °C. Following affinity purification, beads were washed five times with lysis buffer. Purification quality was assessed by running samples on 3–8% Tris-acetate gels (Invitrogen) and western blotting (1% sample, anti-SBP antibody (Sigma-Aldrich)) or silver staining (9% sample, Thermo Scientific).

#### Sample preparation and mass spectrometry by Mortiz group

Samples were resuspended in 100 mM triethylammonium bicarbonate (TEAB) pH 8.5, 2% sodium dodecyl sulfate, boiled at 95 °C for 5 min at 1100 RPM in a thermomixer, and allowed to cool down to room temperature (RT). Tris(2-carboxyethyl)phosphine (TCEP) was added to a final concentration of 5 mM and the mixture was incubated at 60 °C and 1100 RPM for 20 min, allowed to reach RT, iodoacetamide was added to a final concentration of 15 mM and the mixture incubated in the dark at RT and 1100 RPM for 20 min, and TCEP was added to a final total concentration of 10 mM to quench to reaction for 20 min at RT and 1100 RPM.

30 µg each of hydrophilic and hydrophobic carboxylate magnetic SpeedBeads were prepared by three rounds of washing in 500 µL deionized water and then added into the sample. Ethanol (EtOH) was added to a final concentration of ∼70% and incubated for 20min, 1000 RPM at RT. The beads were resolved on a magnet, supernatant was removed, and six rounds of washing in 200 µL 80% EtOH were performed by completely resuspending the beads and then resolving and removing the supernatant. Beads were then resuspended in 100 µL of 50 mM TEAB pH 8.5 and 500 ng of sequencing grade porcine trypsin was added and digestion was allowed to occur overnight at 37 °C, 1000 RPM. Digestion was stopped by the addition of 1% formic acid (FA), the beads were resolved on a magnet, and the supernatant was transferred to fresh microcentrifuge tubes and dried to completion in a SpeedVac and stored at -80 °C until LC-MS analysis.

For LC-MS, samples were resuspended in 10 µL 5% acetonitrile (ACN), 0.1% FA. 9 µL were injected and run. A Neo-Vanquish LC (Thermo) was used to separate peptides on an Easy-Spray PepMap RSLC C18 column (ThermoFisher USA), length 50 cm and inner diameter 75 µm, with a PepMap C18 column (ThermoFisher USA), length 0.5 cm and inner diameter 300 µm with 5 µm dp in-line pre-column, using a 45 min gradient from 0-35% mobile phase B. This was followed by a ramp-up to 80% B over 1 min and held for 15 min as a wash step, then returned to 0% B over 1 min and held for 5 min to re-equilibrate. Mobile phases A and B were 0.1% FA in water and 0.1% FA in ACN. The LC system was attached to a Thermo Orbitrap Lumos. MS measurements were acquired in DDA mode. All samples were analyzed with a spray voltage of 1800 V in positive mode with the RF lens at 30%, using the Orbitrap with an MS1 resolution of 120,000, with normalized automatic gain control = 5e5 and maximum injection time = 50 ms. A cycle time of 3s was specified. MS2 resolution for the Orbitrap was 15,000 (FWHM) at m/z 200, with maximum IT = 22 ms and AGC = 1e5. A normalized collision energy of 30% was used with HCD and an isolation window of 1.6 m/z. Dynamic exclusion was 30 s.

For LC-MS/MS data analysis, Thermo RAW files were converted to mzML format using msConvert (ProteoWizard version 3.0.19225-a1ce12329) ^60^ with “peakPicking true 1-” and “zeroSamples removeExtra” filters. MzML files were searched using Comet ^61^ version 2020.01 rev. 2 with the reviewed SWISS-PROT and TrEMBL *C. elegans* proteome containing 26,706 entries, downloaded May 22, 2023. The database was appended with the green fluorescence protein sequence and the common Repository of Adventitious Proteins (cRAP) database ^62^, and DeBruijn ^63^ randomized decoys (k=2) to the extended database were generated with the Trans-Proteomic Pipeline ^64^ v.6.3.1 (Arcus). The search was performed with a peptide mass tolerance of 20 ppm and a fragment binning tolerance of 0.02, trypsin as the search enzyme and semi-tryptic digestion with two allowed missed cleavages, with carbamidomethyl of cysteines as a fixed modification and methionine oxidation as a variable modification. Comet results were then processed with PeptideProphet ^65^ with accurate mass binning and decoy (DECOY0) hits to pin down the negative distribution with a non-parametric model. The PeptideProphet results were further processed with iProphet ^66^ and protein validation was performed using ProteinProphet ^67^ and quantified at the MS2 level using StPeter ^68^.

#### Sample preparation and mass spectrometry by UF Scripps Proteomics Core

For independent UBR-1 proteomics, separate samples were prepared and evaluated by mass spectrometry at the UF Scripps Proteomics Core. Samples with Dynabeads and purified protein complexes were supplemented with 20% SDS to a final concentration of ∼6% SDS and processed for trypsin digestion using Micro S-Traps^TM^ (Protifi, Huntington, NY) according to manufacturer’s protocol. Briefly, samples were reduced using tris-2(-carboxyethyl)-phosphine (TCEP) at 37°C for 20 minutes, alkylated using methyl methanethiosulfonate (MMTS) at room temperature for 10 minutes, and digested with 1 mg trypsin at 47°C for 1 hour. Subsequently, 40 uL of 50 mM triethylammonium bicarbonate (TEAB) was added and peptides were eluted using centrifugation. Elution was repeated once. A third elution using 35mL of 50% acetonitrile (ACN) was performed. The eluted peptides were then dried under vacuum.

LC-MS/MS analysis of extracted peptides from first replicate set of digests was carried out using an Eclipse Orbitrap Fusion Tribrid mass spectrometer (Thermo Scientific, San Jose, CA), following 2mg capacity ZipTip (Millipore, Billerica, MA) and C18 sample clean-up according to manufacturer’s instructions. Peptides were eluted from an EASY PepMap^TM^ RSLC C18 column (2μm, 100Å, 75μm x 50cm, Thermo Scientific, San Jose, CA) using a gradient of 5-25% solvent B (80/20 acetonitrile/water, 0.1% formic acid) in 135 minutes, followed by 25-44% solvent B in 45 minutes, 44-80% solvent B in 0.10 minute, 10 minute-hold of 80% solvent B, return to 5% solvent B in 3 minutes, and finally another 3-minute hold of 5% solvent B. Gradient was then extended to clean column by increasing solvent B to 98% in 3 minutes, 98% solvent B hold for 10 minutes, return to 5% solvent B in 3 minutes, 5% solvent B fold for 3 minutes, increase of solvent B to 98% in 3 minutes, 98% solvent B hold for 10 minutes, return to 5% solvent B in 3 minutes, 5% solvent B hold for 3 minutes, and finally another increase to 98% solvent B in 3 minutes and a hold of 98% solvent B for 10 minutes. All flow rates were 250nL/minute delivered using a nEasy-LC1000 nano liquid chromatography system (Thermo Scientific, San Jose, CA). Solvent A consisted of water and 0.1% formic acid (FA). Ions were created at 2.3kV using an EASY Spray source (Thermo Scientific, San Jose, CA) held at 50°C. Data dependent scanning was performed by Xcalibur v 4.0.27.10 software using survey scan at 120, 000 resolution in Orbitrap analyzer scanning mass/charge (m/z) 200-2000 followed by higher-energy collisional dissociation (HCD) tandem mass spectrometry (MS/MS) at normalized collision energy of 30% of the most intense ions at maximum speed, at an automatic gain control of 1.0E4. Precursor ions were selected by monoisotopic precursor selection (MIPS) setting to peptide and MS/MS was performed on charged species of 2-8 charges at resolution of 30,000. Dynamic exclusion was set to exclude ions once within 25 second window. All scan events occurred within 2-second specified cycle time. Tandem mass spectra were searched against database made up of proteins downloaded from UniProt for *Caenorhabditis elegans* and *Mus musculus* on July 05, 2023, as well as primary sequence for green fluorescent protein (UniProt Accession P42212). Common contaminant proteins available with Proteome Discoverer v 2.5.0.400 were also used in the search. At the time of the search the combined database with green fluorescent protein contained 81,831 sequences and contaminant protein database contained an additional 298 sequences. All MS/MS spectra were searched using Thermo Proteome Discoverer 2.5.0.400 (Thermo Scientific, San Jose, CA) considering fully tryptic peptides with up to 2 missed cleavage sites. Variable modifications considered during search included methionine oxidation (15.995 Da), asparagine and glutamine deamidation (0.984 Da). Cysteine carbamidomethylation (57.021 Da) was considered as static modification. Proteins were identified at 99% confidence with XCorr score cut-offs offs ^69^ as determined by reverse database search. Protein and peptide identification results were visualized with Scaffold v 5.0.0 (Proteome Software Inc., Portland OR). Scaffold software relies on various search engine results (i.e.: Sequest, X!Tandem, MASCOT) and uses Bayesian statistics to reliably identify spectra ^65^. Proteins were accepted that passed a minimum of two peptides identified at 1% peptide and protein FDR within Scaffold.

LC-MS/MS analysis was repeated with half the amount by volume of extracted peptides using Bruker Tims TOF Pro2 mass spectrometer (Bruker, Bremen, Germany). This was done with second replicate set of digests following ZipTip (Millipore, Billerica, MA) and C18 sample clean up. Tims TOF was coupled to nanoElute ultra-high-performance liquid chromatography (UHPLC) system (plug-in V2.1.60.0; Bruker, Bremen, Germany) equipped with a CaptiveSpray source and a 20μm zero dead volume (ZDV) Sprayer. Peptides were loaded onto Thermo Fisher (Waltham, MA) PepMap Neo C18 trap column (300mm X 5mm, 5mm particle size) and then separated on PepSep Ten Series reverse-phase C18 column (10cm X 75μm, 1.9μm particle size) from Bruker (Bremen, Germany). Column temperature was maintained at 50°C using integrated Bruker Column Toaster (Bremen, Germany). Column was equilibrated using 4 column volumes before loading samples in 100% buffer A (99.9% Fisher Optima® LC/MS water, 0.1% FA) with both steps performed at 800 bar. Trap column was equilibrated at 200 bar. Samples were separated at 500nl/min using linear gradient from 2% to 35% buffer B (99.9% Fisher Optima® LC/MS acetonitrile, 0.1% FA) over 17.8 minutes, ramped to 95% buffer B (in 0.5 minutes), and sustained at 95% buffer B for 2.4 minutes (total separation method time 20.70 min). DDA was performed in PASEF mode with 10 PASEF MS/MS scans. Capillary voltage was set to 1600 V and spectra were acquired in m/z range of 100 to 1700 Th with ion mobility range (1/K0) from 0.7 to 1.5 Vs/cm^2^. Ramp and accumulation time were set to 100 ms to achieve duty cycle close to 100% and total cycle time of 1.17 s. Collision energy was ramped linearly as function of mobility from 65 eV at 1/K0 = 1.6 Vs/cm^2^ to 20 eV at 1/K0 = 0.6 Vs/cm^2^. Precursors with charge state from 0 to 5 (If charge state of peptide precursor is 0, it indicates isotope ion was not detected for that peptide precursor) were selected with target value of 12,500 and intensity threshold of 1000. Any precursors that reached target value in arbitrary units were dynamically excluded for 0.4 min. Tandem mass spectra were searched using MS Fragger ^70^ within Scaffold v 5.0.0 using the same modifications and protein databases as described above.

### Metabolomics

#### Sample Prep

Aqueous metabolites for targeted LC-MS profiling of plasma samples were extracted using a protein precipitation method similar to the one described elsewhere ^71–73^. *C. elegans* samples were first homogenized in 200 µL purified deionized water at 4 _°_C, and then 800 µL of cold methanol containing 124 µM 6C13-glucose and 25.9 µM 2C13-glutamate was added (reference internal standards were added to the samples in order to monitor sample prep). Afterwards samples were vortexed, stored for 30 minutes at -20 _°_C, sonicated in an ice bath for 10 minutes, centrifuged for 15 min at 14,000 rpm and 4_°_C, and then 600 µL of supernatant was collected from each sample (left-over protein pellet was used for BCA assay needed for data normalization). Lastly, recovered supernatants were dried on a SpeedVac (Thermo Fisher) and reconstituted in 0.5 mL of LC-matching solvent containing 17.8 µM 2C13-tyrosine and 39.2 3C13-lactate (reference internal standards were added to the reconstituting solvent in order to monitor LC-MS performance). Samples were transferred into LC vials and placed into a temperature controlled autosampler for LC-MS analysis.

#### LC-MS Assay

Targeted LC-MS metabolite analysis was performed on a duplex-LC-MS system composed of two Shimadzu UPLC pumps, CTC Analytics PAL HTC-xt temperature-controlled auto-sampler and AB Sciex 6500+ Triple Quadrupole MS equipped with an electrospray ionization (ESI) source (1,2,3). UPLC pumps were connected to the autosampler in parallel and could perform two chromatography separations in parallel independently from each other. Each sample was injected twice on two identical analytical columns (Waters XBridge BEH Amide Premier 2.1 x 150mm, Waters part # 186009930) performing separations in hydrophilic interaction liquid chromatography (HILIC) mode. While one column performed separation for MS data acquisition in ESI+ ionization mode, the other column was equilibrated for sample injection, chromatography separation and MS data acquisition in ESI-mode. Each chromatography separation was 16 min (total analysis time per sample is 32 min), and MS data acquisition was performed in multiple-reaction-monitoring (MRM) mode. The LC-MS system was controlled using AB Sciex Analyst 1.7.1 software. The LC-MS assay targeted 371 metabolic species and 32 SILISs (403 MRMs). In addition to the study samples, two sets of quality control (QC) samples were used to monitor the assay performance and data reproducibility. One in-house QC [QC(I)] was a pooled human serum sample used to monitor system performance, and the other QC [QC(S)] consisted of pooled study samples, which was used to monitor data reproducibility. Each QC sample was injected once for every 10 study samples. Two sets of samples, six samples each, were analyzed in two separate batches. 243 metabolites (plus four spiked standards) were measured across the study set and the median CV was 2.1 %for the first set and 3.4 % for the second set (based on 2 QC injections per batch and measured peak areas without any signal normalization). Metabolites were relatively quantified as area-under-the-curves of the MRM transitions using AB Sciex MultiQuant 3.0.3 software. The data were normalized using the total protein count (BCA assay) of each sample.

### Statistical analysis

#### High-intensity locomotor behavior and liquid aldicarb assays

For MWT assays, high-intensity locomotor behavior was recorded on plates with 4-5 wells per genotype and 4 animals per well. Data were collected from at least 3 independent biological replicates from separate parent plates. In line plots, solid lines represent the average speed of all animals and shaded regions are SEM. For aldicarb assays, data for each genotype was normalized to the average speed of a 10 min baseline without drug. Bar graphs represent the average of all wells for indicated time point and genotype, dots represent single wells tracked (4 animals/well), and error bars are SEM. Significance for data shown in line plots were tested with pairwise two-way ANOVA. Significance for bar and dot graphs was tested using one-way ANOVA and Bonferroni’s *post-hoc* correction for multiple comparisons. Significance was defined as p<0.05. Statistical analysis was performed using GraphPad Prism software.

#### Developmental viability assays

Each dataset was composed of 9-10 total plates for each genotype. Data was derived from 3 independent experiments with 3-4 plates per experiment for each genotype. Statistical significance was determined using one-way ANOVA with Bonferroni’s *post-hoc* correction for multiple comparisons.

#### Proteomics

For AP-proteomics, proteins were only considered hits if they were identified with 1.5x or greater enrichment in GFP::UBR-1 test samples over GFP::SL2::UBR-1 negative control samples. Significance was tested using unpaired, nonparametric Mann-Whitney tests with normalized total spectral counts across ten independent proteomic experiments (See Supp data). These comparisons are reported as p values. To correct for multiple comparisons, we applied a 5% false discovery rate (FDR) and a *post hoc* Benjamini-Hochberg method to Mann-Whitney tests. These comparisons are reported as q values. Each sample preparation and its corresponding mass spectrometry experiment was considered an individual n for analysis. For statistical comparisons in Fig 3, we normalized total spectral counts for the candidate hit protein to both its molecular weight as well as the total spectral counts and molecular weight of the affinity purification target, as previously detailed ^35^. Statistical analysis was performed using GraphPad Prism software.

### Data availability

Strains and plasmids are available upon request. The authors affirm that all data necessary for confirming the conclusions of the article are present within the article, figures, tables and supplementary materials.

Metadata (excel) and statistical analysis (prism) for all quantitative behavioral and viability experiments can be found with the following DOI (10.5061/dryad.8cz8w9h3r) on the Dryad website (https://datadryad.org).

Mass spectrometry files for UBR-1 AP-proteomics and SL2 negative controls have been uploaded to the PRIDE database (https://www.ebi.ac.uk/pride/) as Project number: PXD066115.

## Acknowledgements

We would like to thank Dr. George Tsaprailis and the UF Scripps Proteomics Core for independent mass spectrometry. We thank Daniel Raftery, Danijel Djukovic, Wentao Zhu and the entire staff of the Northwest Metabolomics Research Center at the University of Washington for performing LC-MS for glutamate metabolite data acquisition and analysis training. We are grateful to Research Scientific Computing at SCRI for providing HPC resources for processing AlphaFold predictions. We thank Dr. Chris Rongo for generously providing *odIs6*. We thank SHIGEN for *tm5996*, the WormBase genetic resource database, and Alliance of Genome Resources. B.G. and N.Z. were supported by National Institutes of Health Grant R01 DA056370. J.P. was supported by NIH Fellowship F32GM156113. N.Z. is a Howard Hughes Medical Institute investigator. This work was supported in part by the National Institutes of Health grants R01 GM087221 (R.L.M.), S10 OD026936 (R.L.M.), and by a National Science Foundation grant MRI-1920268 (R.L.M.).

## Author Contributions

J.S.P performed *C. elegans* genetics, data acquisition, and data analysis for all behavioral assays, expression studies, AlphaFold predictions. J.S.P. performed *C. elegans* culture, protein extraction and protein purification for proteomics. J.S.P. and K.J.O. created all CRISPR engineered, deletion mutant and transgenic rescue strains. J.S.P. performed glutamate metabolite data analysis. R.L.M., S.M. and M.M. performed LC-MS/MS. R.L.M., S.M., M.M., C.K. and J.S.P. analyzed mass spectrometry data. N.D.M. provided conceptual and technical input. B.G. provided input, advised on experimental design, and provided supervision for personnel. N.Z. and D.P. provided conceptual input. B.G., N.Z. and R.L.M. managed project funding. J.S.P. and B.G. wrote the manuscript. R.M, N.Z. and D.P. edited the manuscript.

---

**Supplementary Fig 1:**
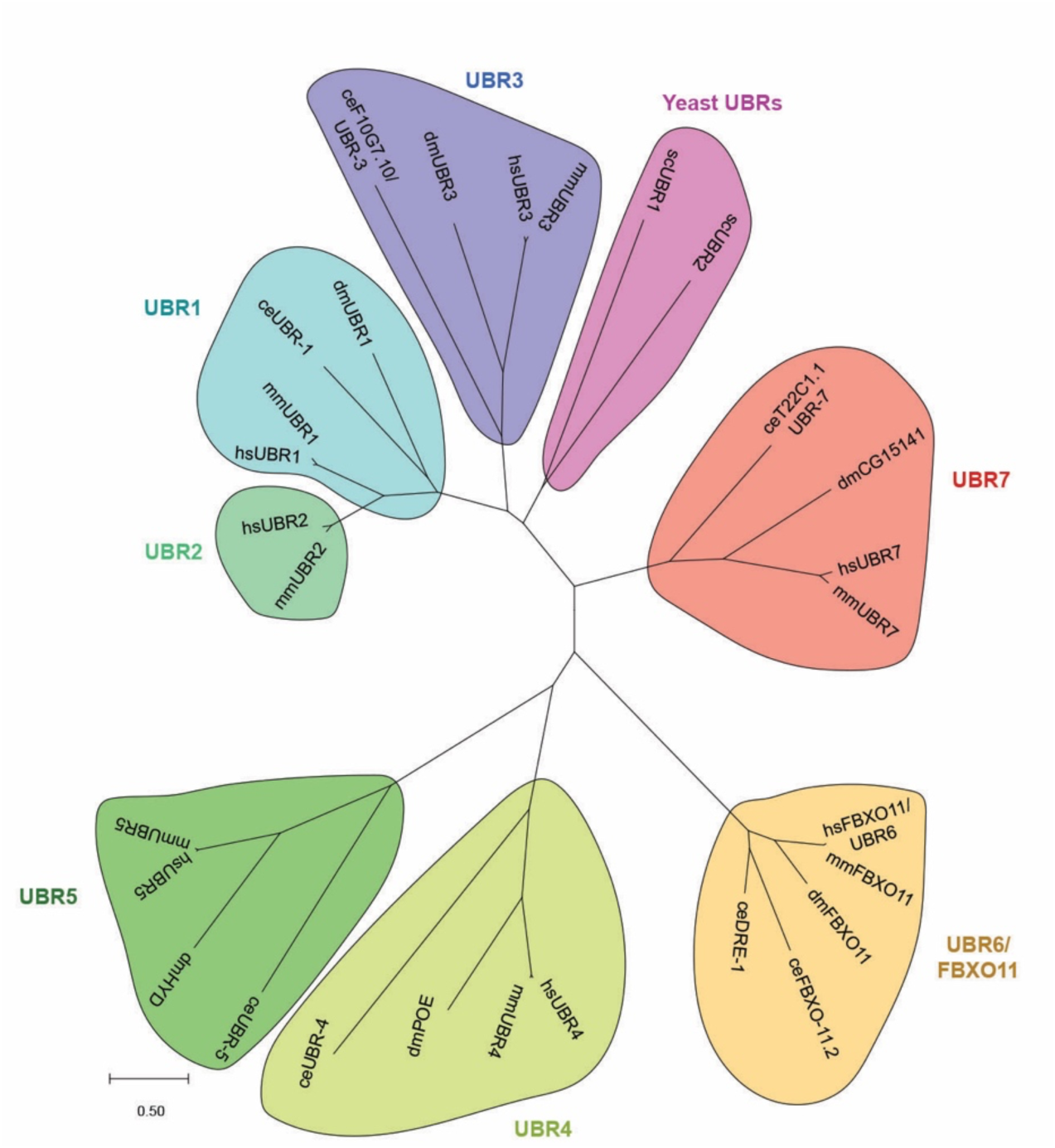
UBR family proteins are conserved across phylogenetic clades from yeast through mammals. Phylogenetic tree shows UBR protein family, including *C. elegans* UBR-1 and its mammalian orthologs UBR1 and UBR2, are conserved from single-celled eukaryotes through vertebrates. Protein sequences from yeast (*Saccharomyces cerevisiae*, sc), *C. elegans* (ce), fruitfly (*Drosophila melanogaster*, dm), mouse (*Mus musculus*, mm), and human (*Homo sapiens*, hs) were aligned using MUSCLE and built into a maximum likelihood phylogenetic tree using MEGA11 maximum likelihood analysis method.

**Supplementary Fig 2:**
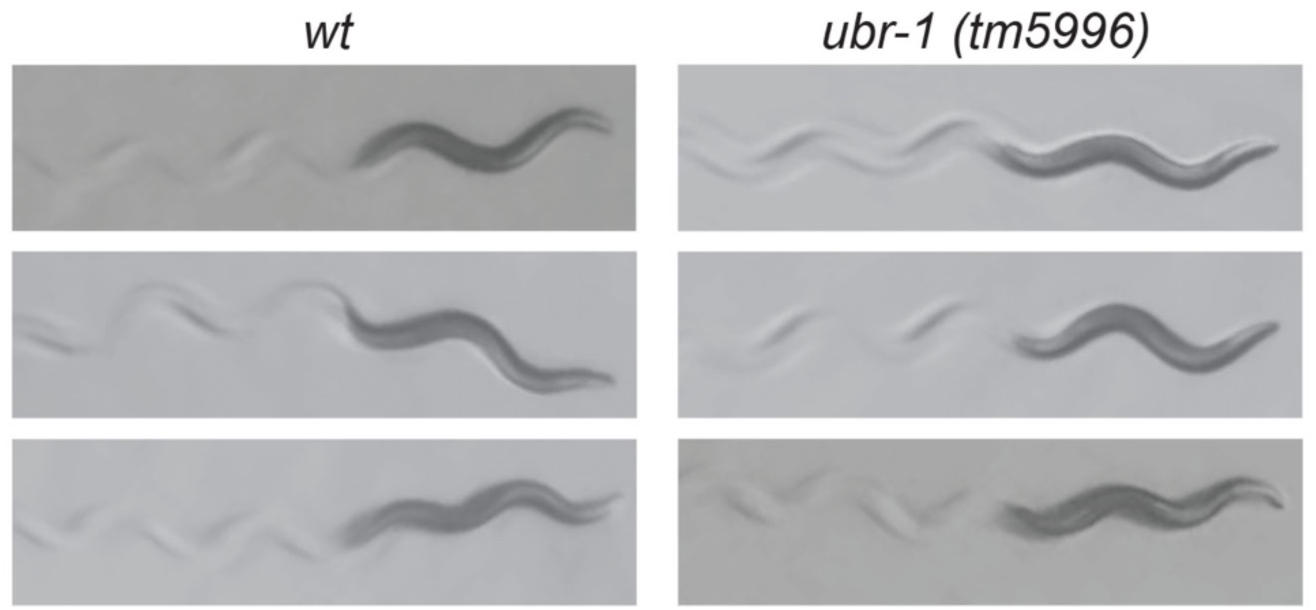
*ubr-1* mutants do not display forward locomotion defects during low-intensity locomotion on solid media. Still frame images showing *C. elegans* and tracks on solid media plates for indicated genotypes. Note *ubr-1(tm5996)* mutants display similar forward locomotion to wild type on solid media. Animals were placed on a solid-media growth plate and allowed to crawl for 1 min before recording.

**Supplementary Fig 3:**
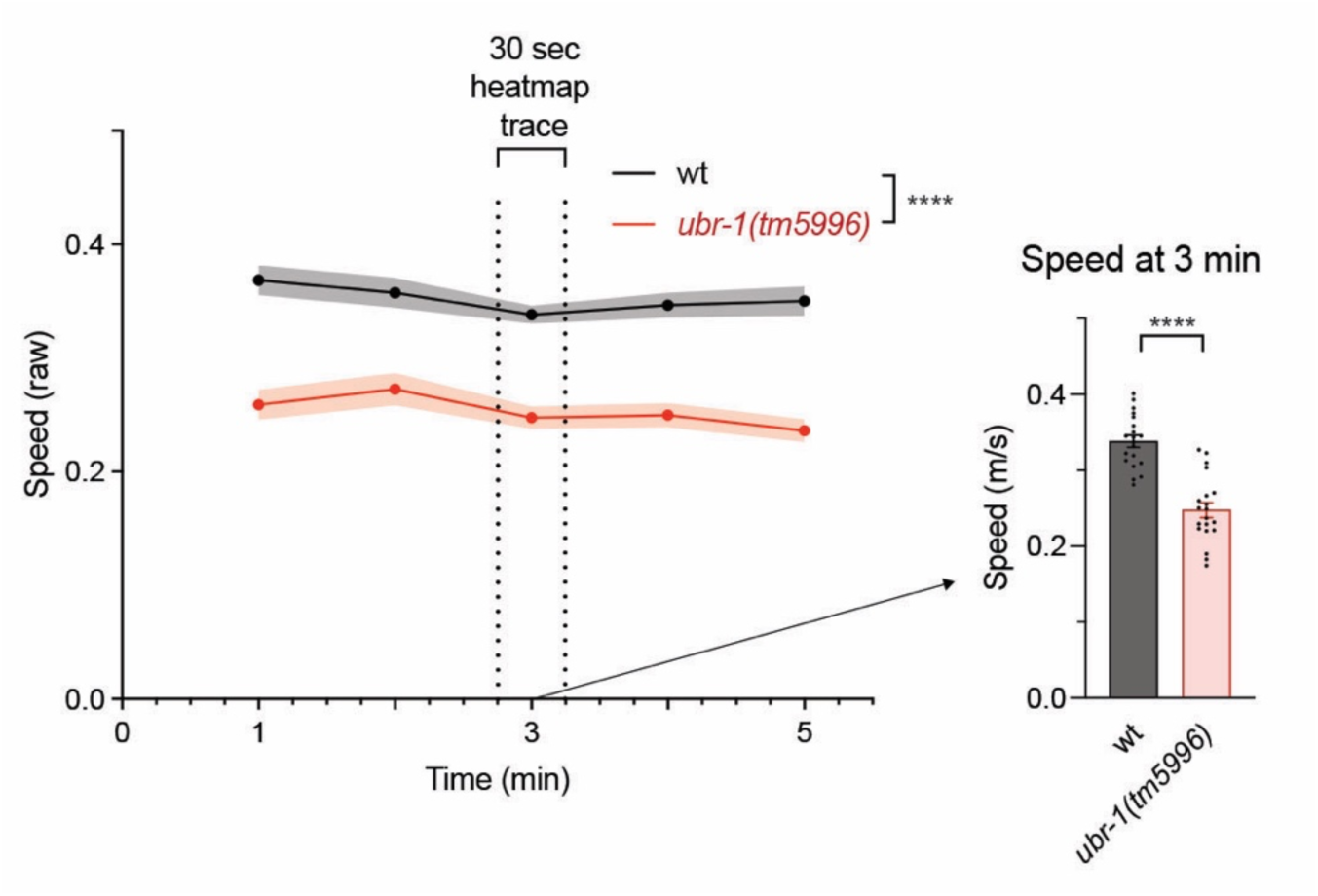
Expanded analysis of UBR-1 effects on high-intensity locomotor speed at early time points. Quantitation of high-intensity swimming locomotor speeds for wild type and *ubr-1* mutants across time window used to create MWT traces of individual animals shown in Figure 1D. 30 sec Heatmap trace time window for Figure 1D is annotated (between dashed lines). Shown are MWT plots of average locomotor speed (left) and expanded quantitation at 3 minutes (right). MWT plots (solid lines, left) represent average speed of all recorded animals (4 animals/well, 5 wells per genotype per experiment, and 3-5 independent experiments) and shaded regions are SEM. Bars represent average of all wells for indicated time point, dots represent single wells tracked (4 animals/well), and error bars are SEM. Significance for plots (genotype annotations, left) tested with pairwise two-way ANOVA, and significance for bars with dots (right) tested using an unpaired two-tailed Student’s *t*-test. **** p<0.0001

**Supplementary Fig 4:**
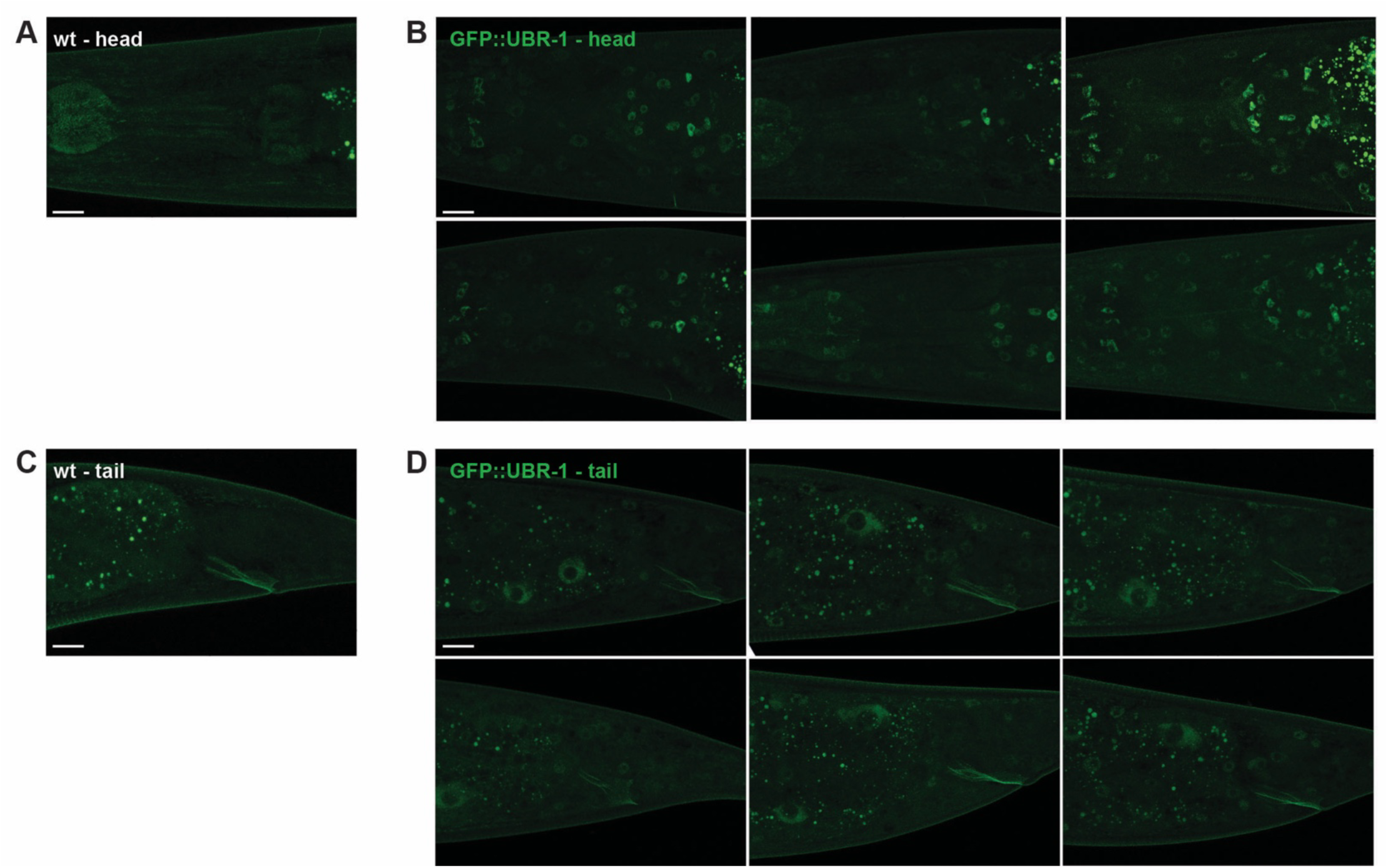
Expanded dataset showing CRISPR engineered GFP::UBR-1 expression in head and tail neurons. **A**. Super-resolution image of adult *C. elegans* head. In wild type, autofluorescence shows animal’s body outline, pharynx, and gut granules (bright puncta far right). **B**. Further examples of super-resolution images showing GFP::UBR-1 expression in head neurons, continued from Figure 2C. Note GFP::UBR-1 is excluded for neuronal nuclei. **C**. Super-resolution image of tail in wild type where autofluorescence shows animal’s body outline, anus and gut granules (bright puncta left). **D**. Examples of Super-resolution images showing GFP::UBR-1 expression in tail neurons. Scale bar is 10 µm

**Supplementary Fig 5:**
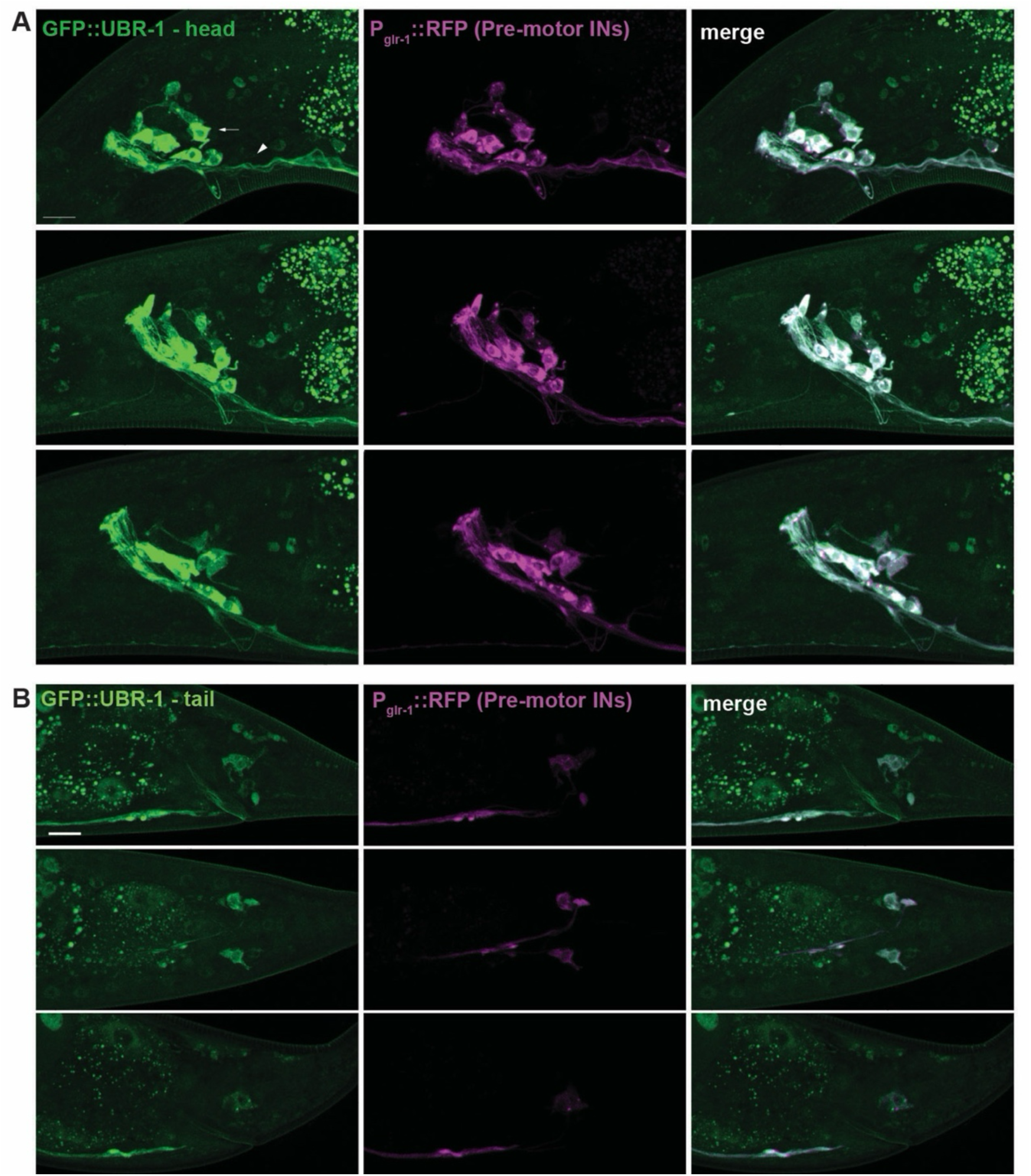
Expanded dataset showing CRISPR engineered GFP::UBR-1 expression in pre-motor interneurons. **A and B**. Super-resolution images of GFP::UBR-1 (green) localized to cell bodies (arrows) and axons (arrowhead) of pre-motor interneurons in head (**A**) and tail (**B**). Note GFP::UBR-1 is excluded for neuronal nuclei. Interneurons visualized using P*glr-1*::RFP (magenta). Scale bar is 10 µm.

**Supplementary Fig 6:**
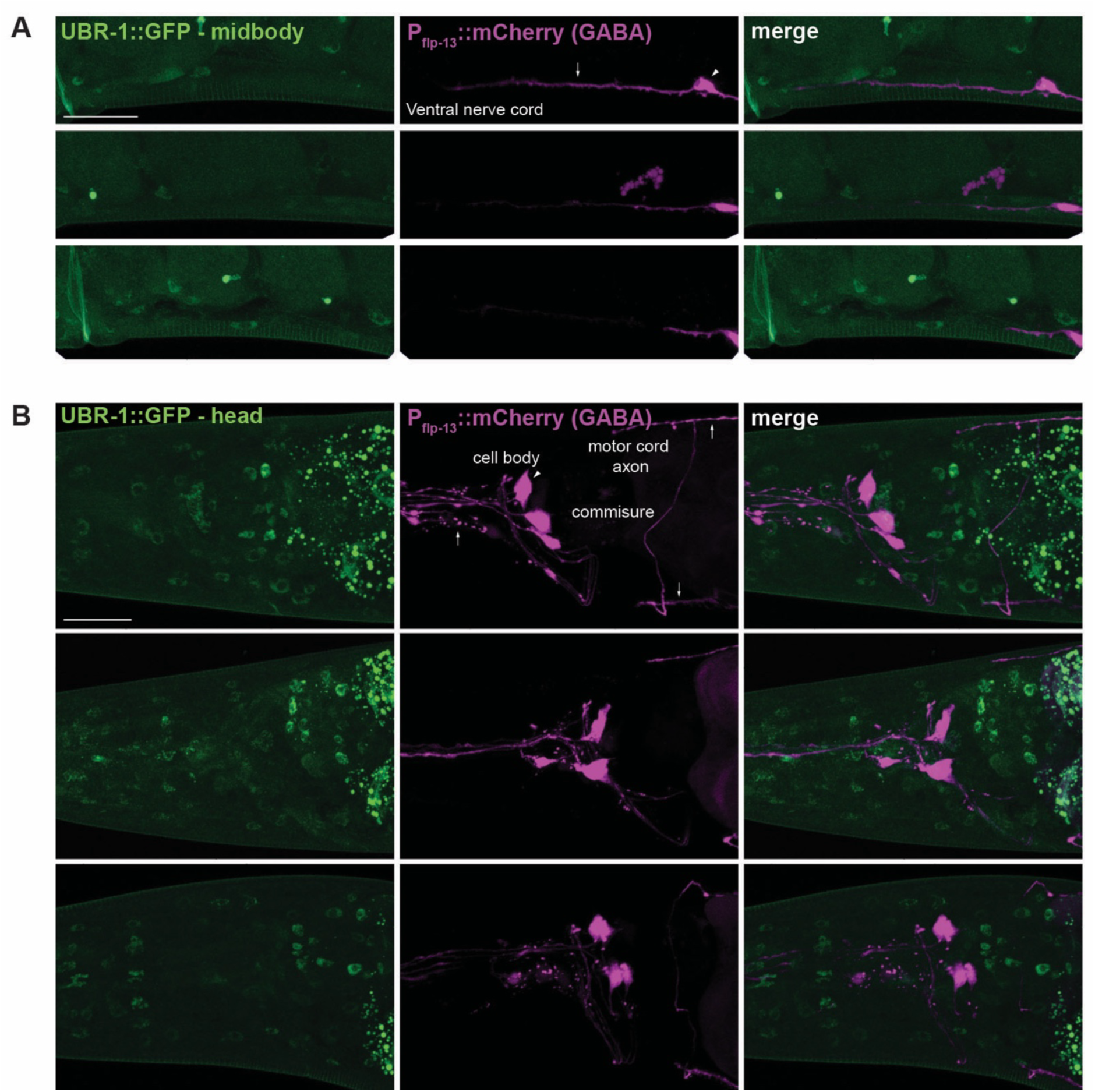
CRISPR engineered GFP::UBR-1 is not expressed in GABA motor neurons. **A-B**. Super-resolution images showing GFP::UBR-1 (green) does not localize to cell bodies (arrows) or axons (arrowheads) of inhibitory GABA DD motor neurons in ventral cord (**A**) or GABA neurons in head (**B**). GABA neurons visualized using P*_flp-13_*::mCherry (magenta). Scale bar is 10 µm.

**Supplementary Fig 7:**
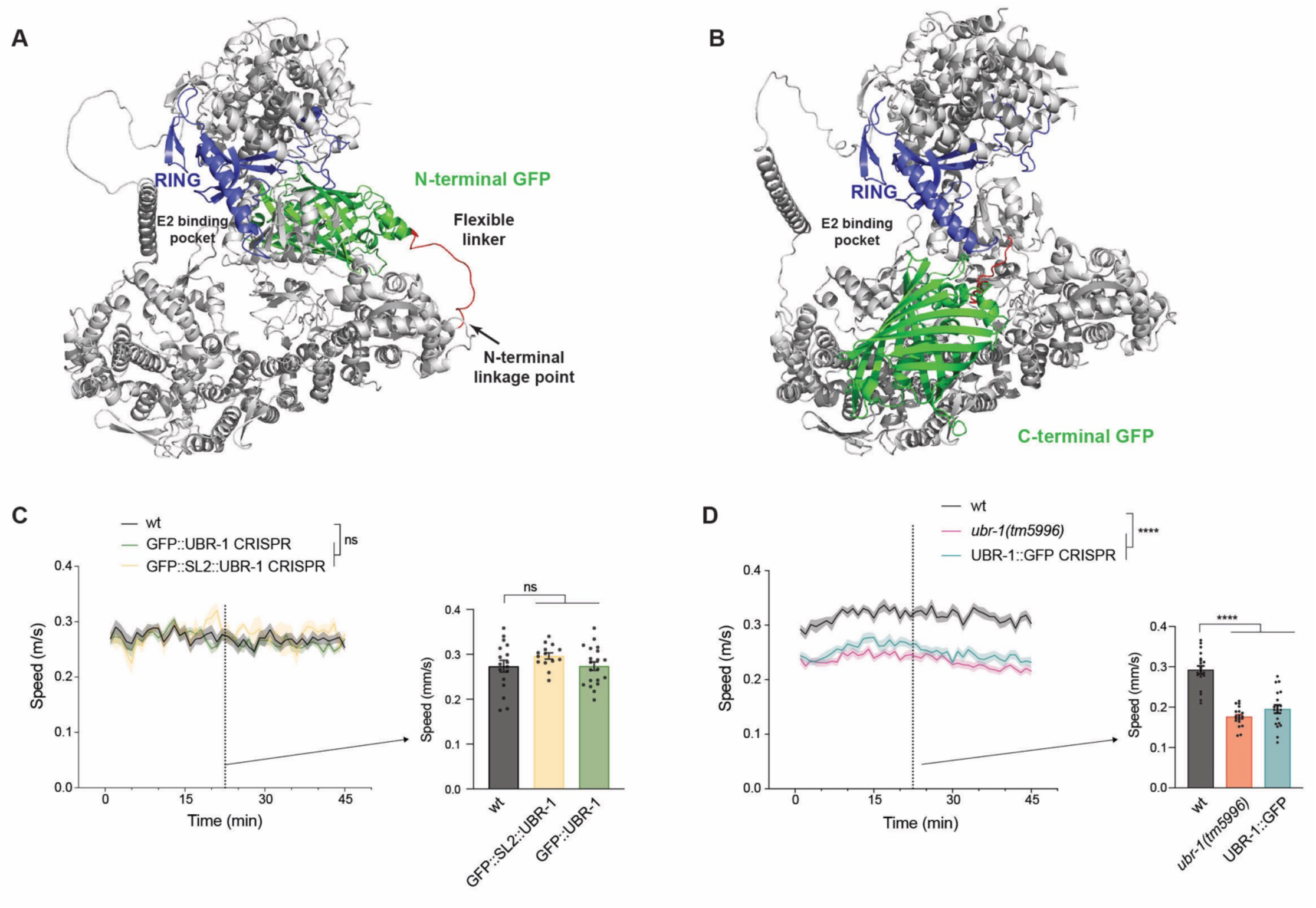
Evaluating CRISPR engineered GFP tag location on UBR-1 function. **A and B**: AlphaFold predicted structures of N-terminal-tagged GFP::UBR-1 (**A**) and C-terminal-tagged UBR-1::GFP (**B**). Highlighted are GFP (green), catalytic RING domain (blue) and flexible linker (red). N-terminal GFP is away from RING domain. C-terminal GFP is positioned near catalytic RING suggesting it could sterically hinder binding of E2 or ubiquitin that is transferred to substrate. Note disordered regions were removed from structural prediction. **C**. Quantitation shows locomotor speed is not significantly altered in GFP::UBR-1 CRISPR engineered animals compared to GFP::SL2::UBR-1 negative control or wild type. Shown are MWT plots of average locomotor speed (left) and expanded quantitation at 23 minutes (right). **D**. Quantitation shows UBR-1::GFP CRISPR engineered animals have impaired locomotor speed similar to *ubr-1* mutants. **For C and D**, MWT plots (solid lines, left) represent average speed of all recorded animals (4 animals/well, 5 wells per genotype per experiment, and 3-5 independent experiments) and shaded regions are SEM. Bars represent average of all wells for indicated time point, dots represent single wells tracked (4 animals/well), and error bars are SEM. Significance for plots (genotype annotations, left) tested with pairwise two-way ANOVA, and significance for bars with dots (right) tested using one-way ANOVA and Bonferroni’s post-hoc correction. **** p<0.0001, ns = not significant

**Supplementary Fig 8:**
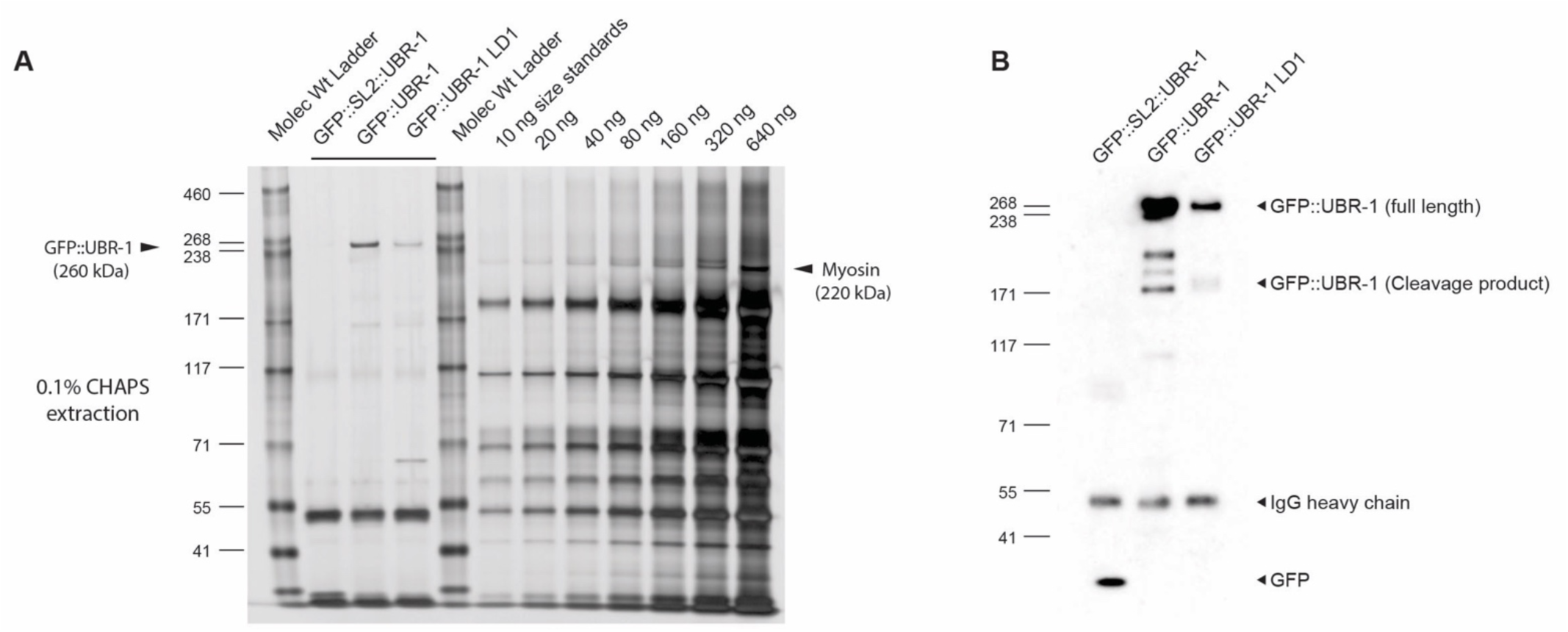
Expanded dataset for quality control of *C. elegans* proteomics. **A and B**. Further examples of silver stain (**A**) and immunoblot (**B**) of anti-GFP precipitated samples used to evaluate sample quality prior to proteomics for GFP::UBR-1 and GFP::SL2::UBR-1 (negative control). Titrated Myosin standards were used to quantify amount of GFP::UBR-1 to ensure sufficient purification target was present for proteomics. Note GFP::UBR-1 LD1 construct was also evaluated in this preparation.

**Supplementary Fig 9:**
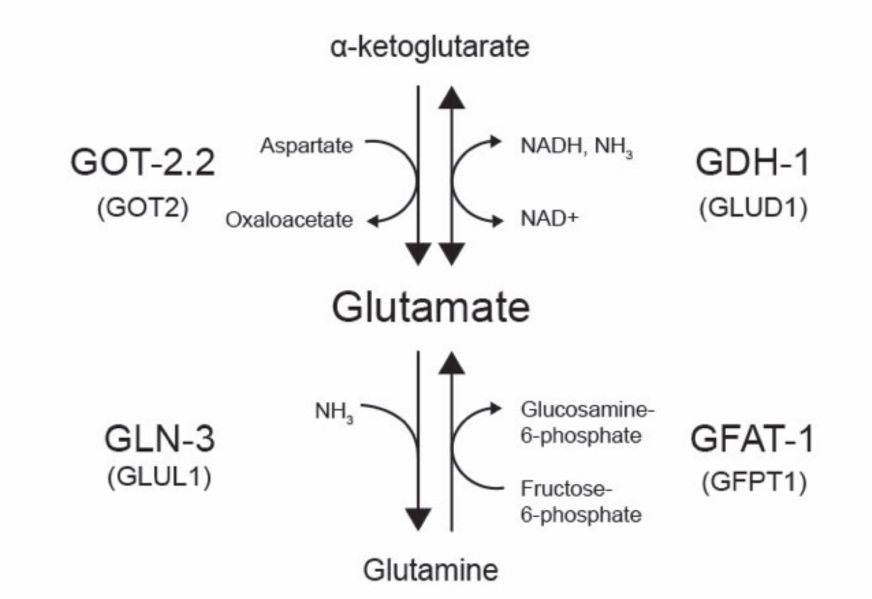
Overview of GOT-2.2, GLN-3, GFAT-1 and GDH-1 effects on glutamate metabolism in *C. elegans*. Schematic of glutamate metabolism showing GOT-2.2 and GFAT-1 promote generation of glutamate. GLN-3 has an opposing role converting glutamate to glutamine. GDH-1 catalyzes reversible reaction of glutamate to ɑ-ketoglutarate.

**Supplementary Fig 10:**
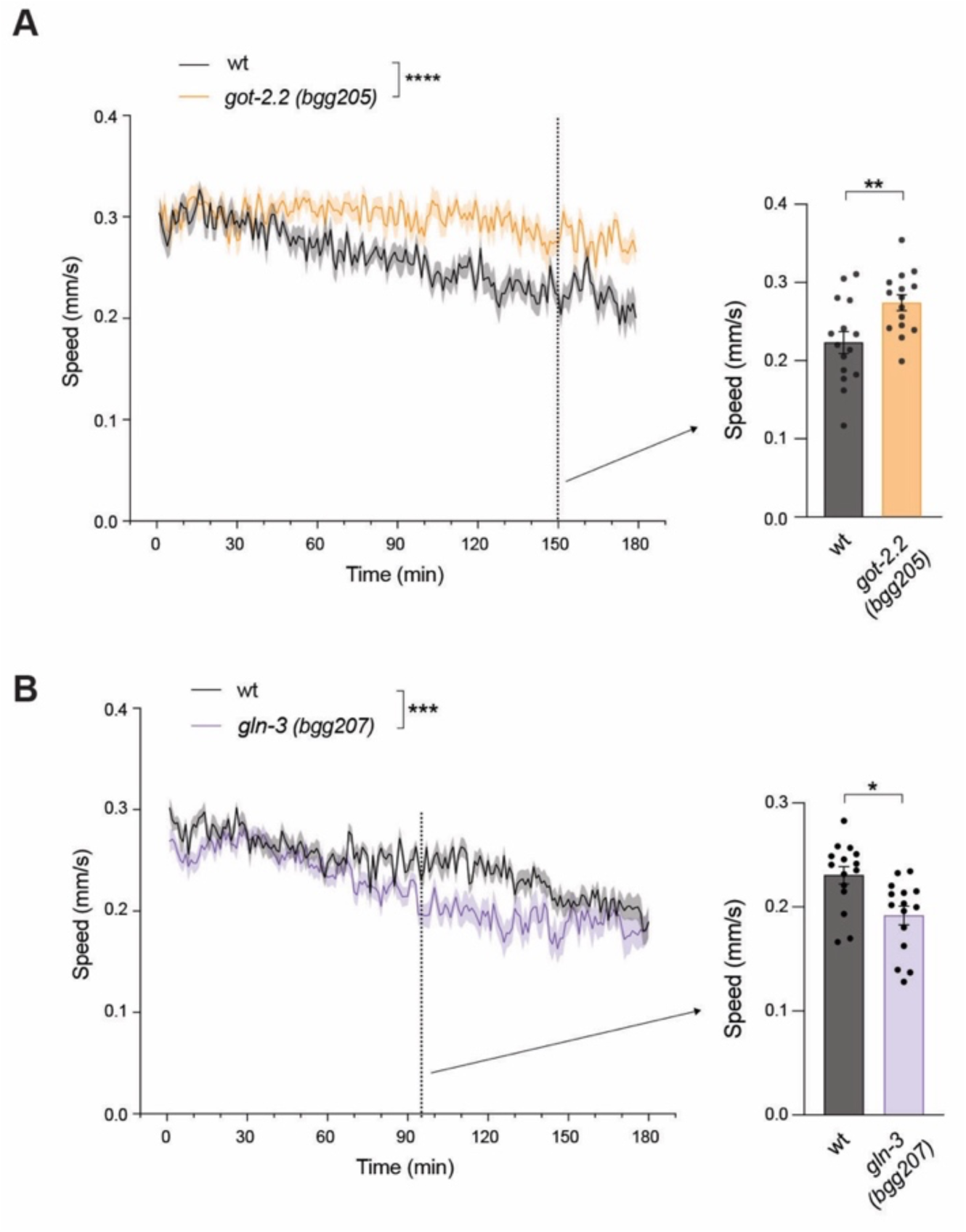
Expanded dataset showing further mutant alleles of *got-2* and *gln-3* affect high-intensity locomotor activity. **A**. Quantitation of swimming speed shows sustained locomotion in second mutant allele of *got-2.2 (bgg205)* compared to wild type. Shown are MWT plots of average locomotor speed (left) and expanded quantitation at 150 mins (right). **B**. Quantitation shows second mutant allele of *gln-3 (bgg207)* displays early fatigue compared to wild type. Shown are MWT plots of average locomotor speed (left) and expanded quantitation at 95 mins (right). **For A and B**, MWT plots (solid lines, left) represent average speed of all recorded animals (4 animals/well, 5 wells per genotype per experiment, and 3 independent experiments) and shaded regions are SEM. Bars represent average of all wells for indicated time point, dots represent single wells tracked (4 animals/well), and error bars are SEM. Significance for plots (genotype annotations, left) tested with pairwise two-way ANOVA, and significance for bars with dots (right) tested using unpaired two-tailed Student’s *t*-test. **** p<0.0001, *** p<0.001, ** p<0.01, * p<0.05

**Supplementary Fig 11:**
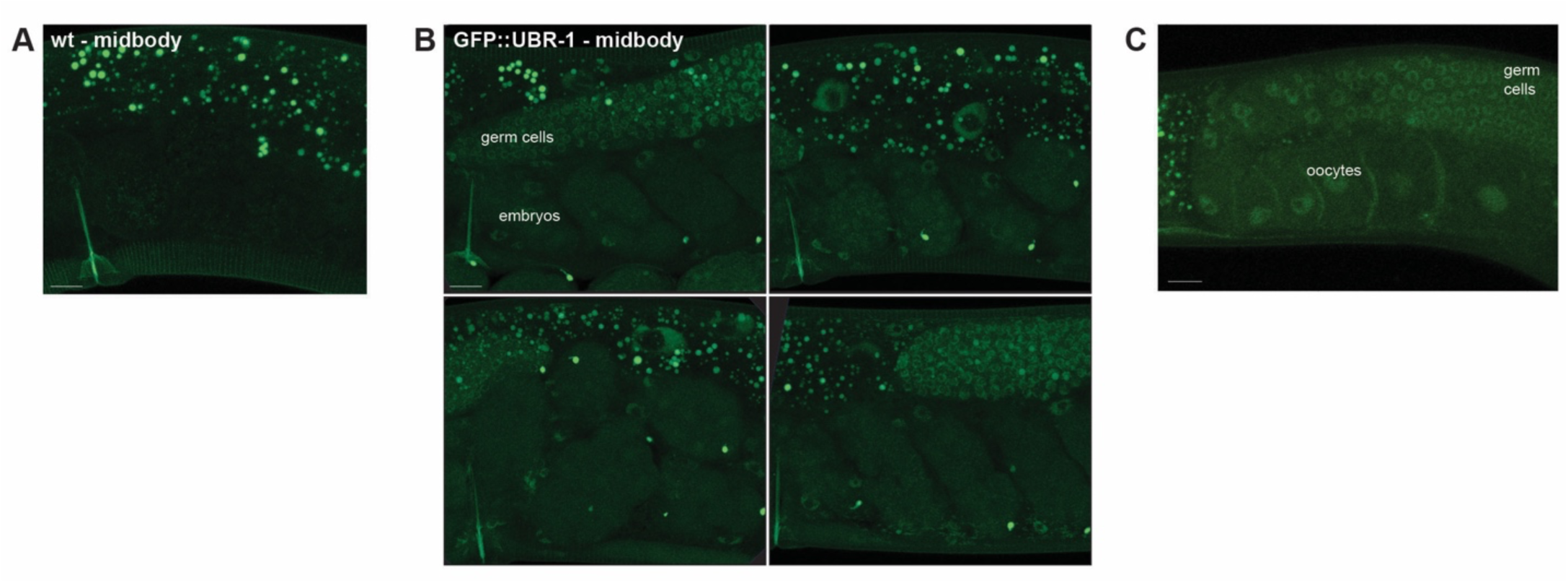
Expanded dataset showing CRISPR engineered GFP::UBR-1 expression in gonad. **A**. Super-resolution image of adult *C. elegans* midbody. In wild type, autofluorescence shows body outline, vulva and gut granules (bright puncta far right). **B**. Further examples of super-resolution images showing GFP::UBR-1 expression in germ cells and embryos. **C**. Further examples of GFP::UBR-1 expression in oocytes and germ cells. **For B and C**, images are continued examples from Figure 6B and C. Scale bar is 10 µm.

**Supplementary Fig 12:**
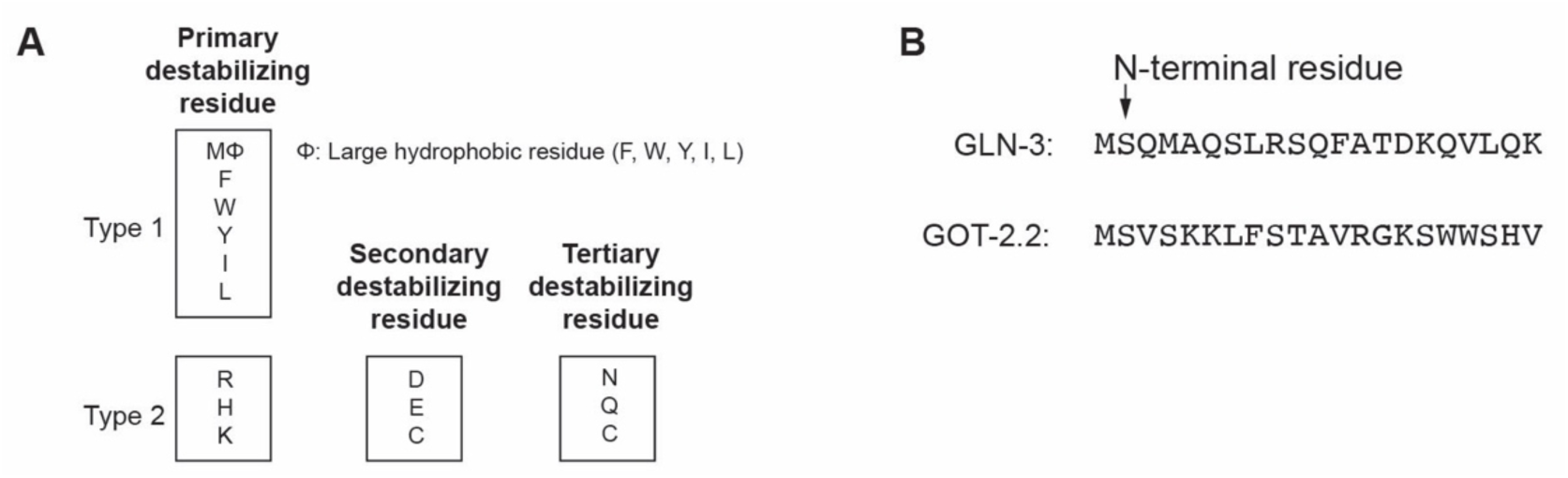
GLN-3 and GOT-2.2 do not have predicted UBR-1 substrate recognition degron sequences. **A**. Diagram of known primary, secondary, and tertiary N-terminal destabilizing residues (N-degrons) recognized by UBR-box proteins to trigger N-end rule ubiquitination and degradation. **B**. N-terminal amino acid sequence for GLN-3 and GOT-2.2. Note first residues following initial methionine are not classified as N-degrons for both proteins.

**Supplementary Table 1:**
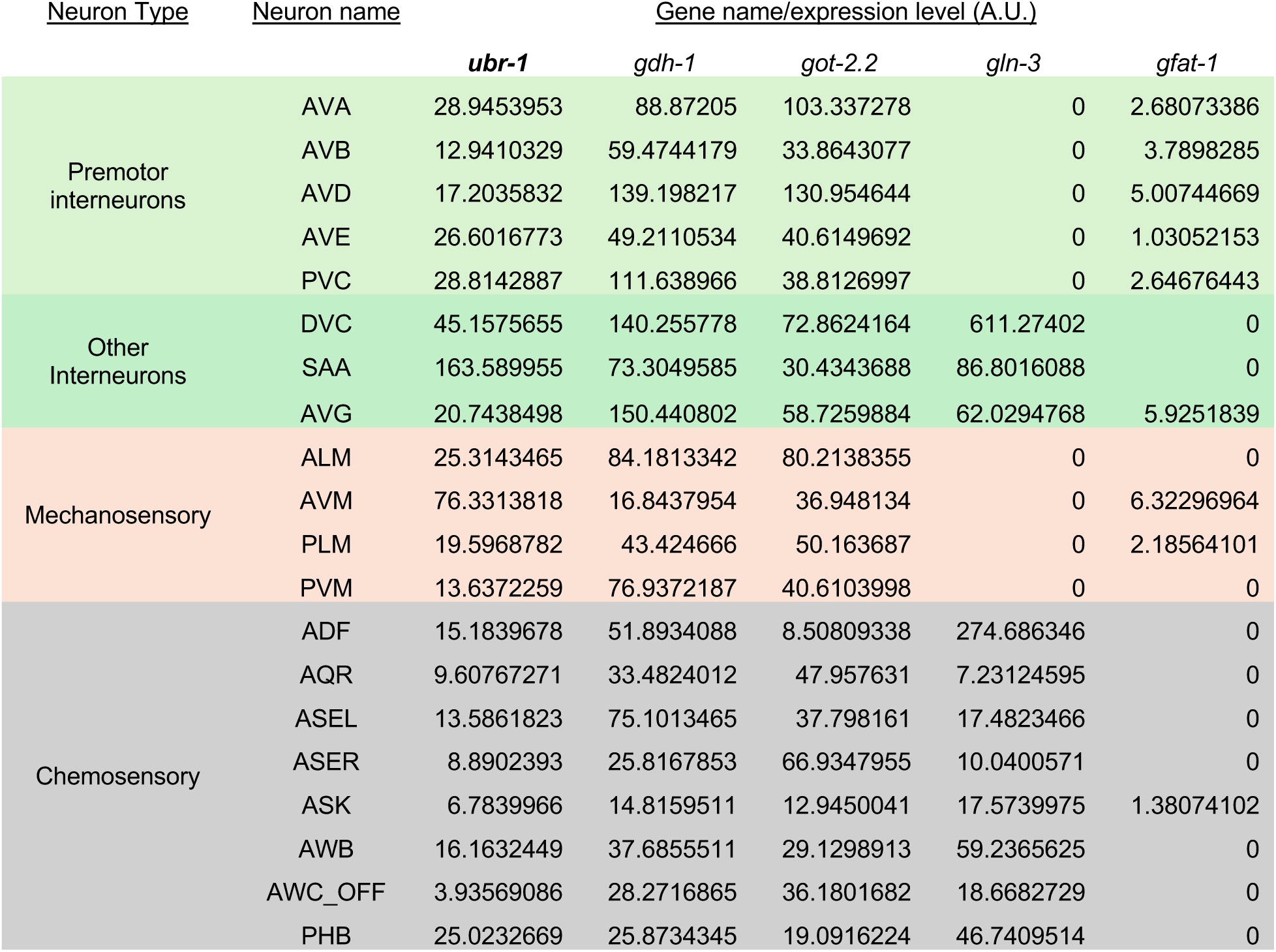
CeNGEN transcriptional profiles for *ubr-1* and glutamate metabolic enzymes. CeNGEN single cell neural transcriptional atlas for *C. elegans* was analyzed and expression of *ubr-1*, *gdh-1*, *got-2.2*, *gln-3*, and *gfat-1* were found in pre-motor interneurons, other interneurons and sensory neurons.

| Neuron Type | Neuron name | Gene name/expression level (A.U.) |  |  |  |  |
| --- | --- | --- | --- | --- | --- | --- |
|  |  | *ubr-1* | *gdh-1* | *got-2.2* | *gln-3* | *gfat-1* |
| Premotor interneurons | AVA | 28.9453953 | 88.87205 | 103.337278 | 0 | 2.68073386 |
|  | AVB | 12.9410329 | 59.4744179 | 33.8643077 | 0 | 3.7898285 |
|  | AVD | 17.2035832 | 139.198217 | 130.954644 | 0 | 5.00744669 |
|  | AVE | 26.6016773 | 49.2110534 | 40.6149692 | 0 | 1.03052153 |
|  | PVC | 28.8142887 | 111.638966 | 38.8126997 | 0 | 2.64676443 |
| Other Interneurons | DVC | 45.1575655 | 140.255778 | 72.8624164 | 611.27402 | 0 |
|  | SAA | 163.589955 | 73.3049585 | 30.4343688 | 86.8016088 | 0 |
|  | AVG | 20.7438498 | 150.440802 | 58.7259884 | 62.0294768 | 5.9251839 |
| Mechanosensory | ALM | 25.3143465 | 84.1813342 | 80.2138355 | 0 | 0 |
|  | AVM | 76.3313818 | 16.8437954 | 36.948134 | 0 | 6.32296964 |
|  | PLM | 19.5968782 | 43.424666 | 50.163687 | 0 | 2.18564101 |
|  | PVM | 13.6372259 | 76.9372187 | 40.6103998 | 0 | 0 |
| Chemosensory | ADF | 15.1839678 | 51.8934088 | 8.50809338 | 274.686346 | 0 |
|  | AQR | 9.60767271 | 33.4824012 | 47.957631 | 7.23124595 | 0 |
|  | ASEL | 13.5861823 | 75.1013465 | 37.798161 | 17.4823466 | 0 |
|  | ASER | 8.8902393 | 25.8167853 | 66.9347955 | 10.0400571 | 0 |
|  | ASK | 6.7839966 | 14.8159511 | 12.9450041 | 17.5739975 | 1.38074102 |
|  | AWB | 16.1632449 | 37.6855511 | 29.1298913 | 59.2365625 | 0 |
|  | AWC_OFF | 3.93569086 | 28.2716865 | 36.1801682 | 18.6682729 | 0 |
|  | PHB | 25.0232669 | 25.8734345 | 19.0916224 | 46.7409514 | 0 |

**Supplementary Table 2:**
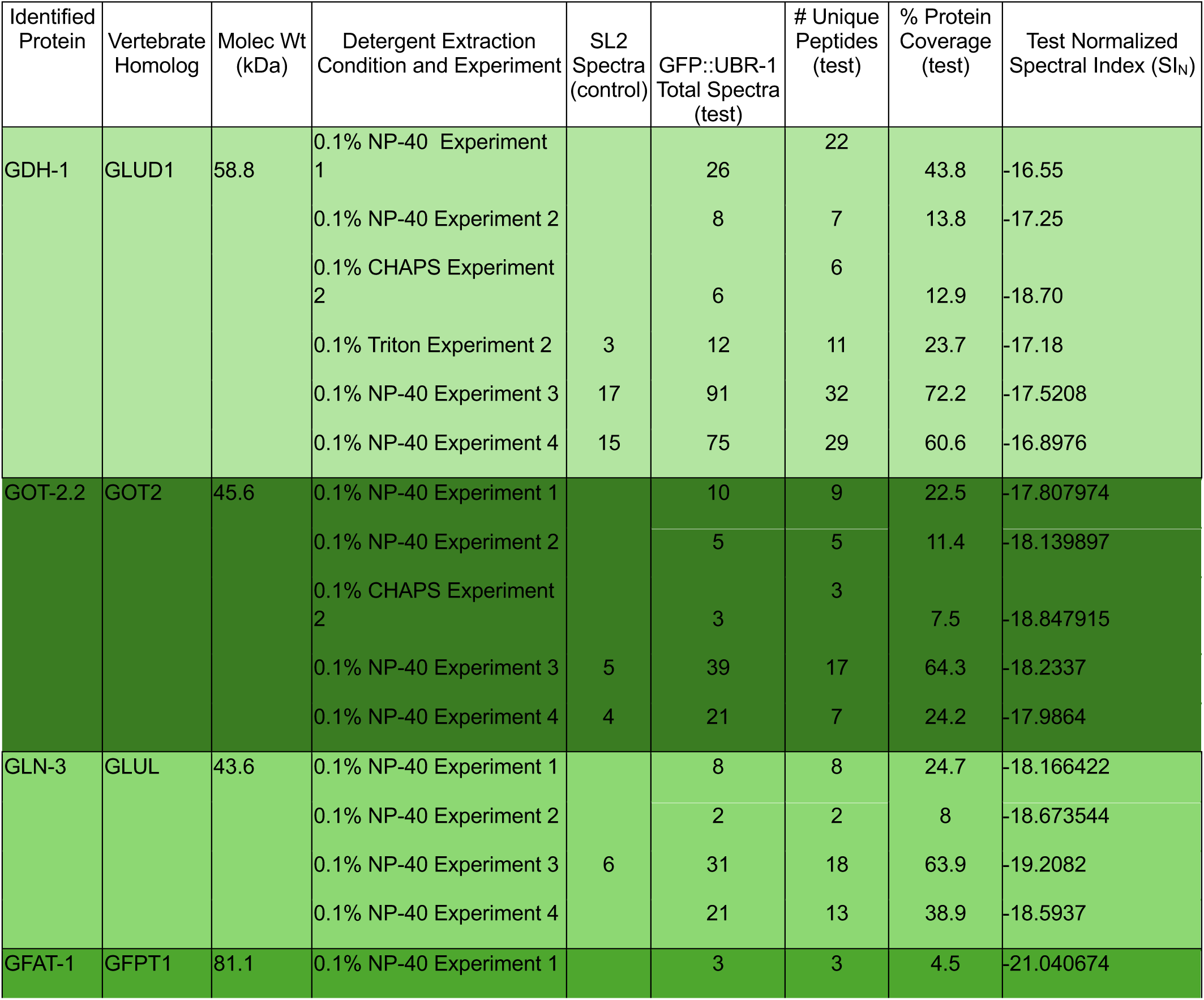
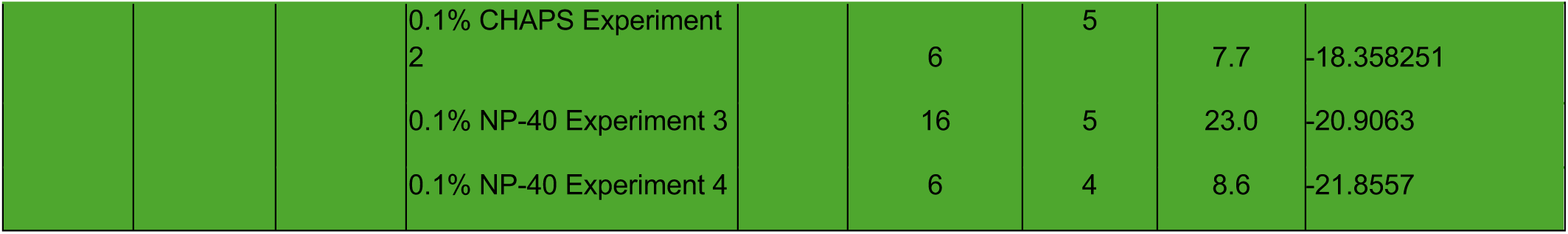
Expanded dataset for UBR-1 proteomic hits. Mass spectrum data are presented for each individual experiment where glutamate metabolic enzymes were identified as UBR-1 proteomic hits. Shown is percent coverage of protein sequences for individual hits in GFP::UBR-1 test samples. Normalized spectral index (SI_N_) for each test sample was calculated using StPeter as a label-free quantitative measure of relative abundance for protein hits where smaller values represent higher abundance.

**Supplementary Table 3:**
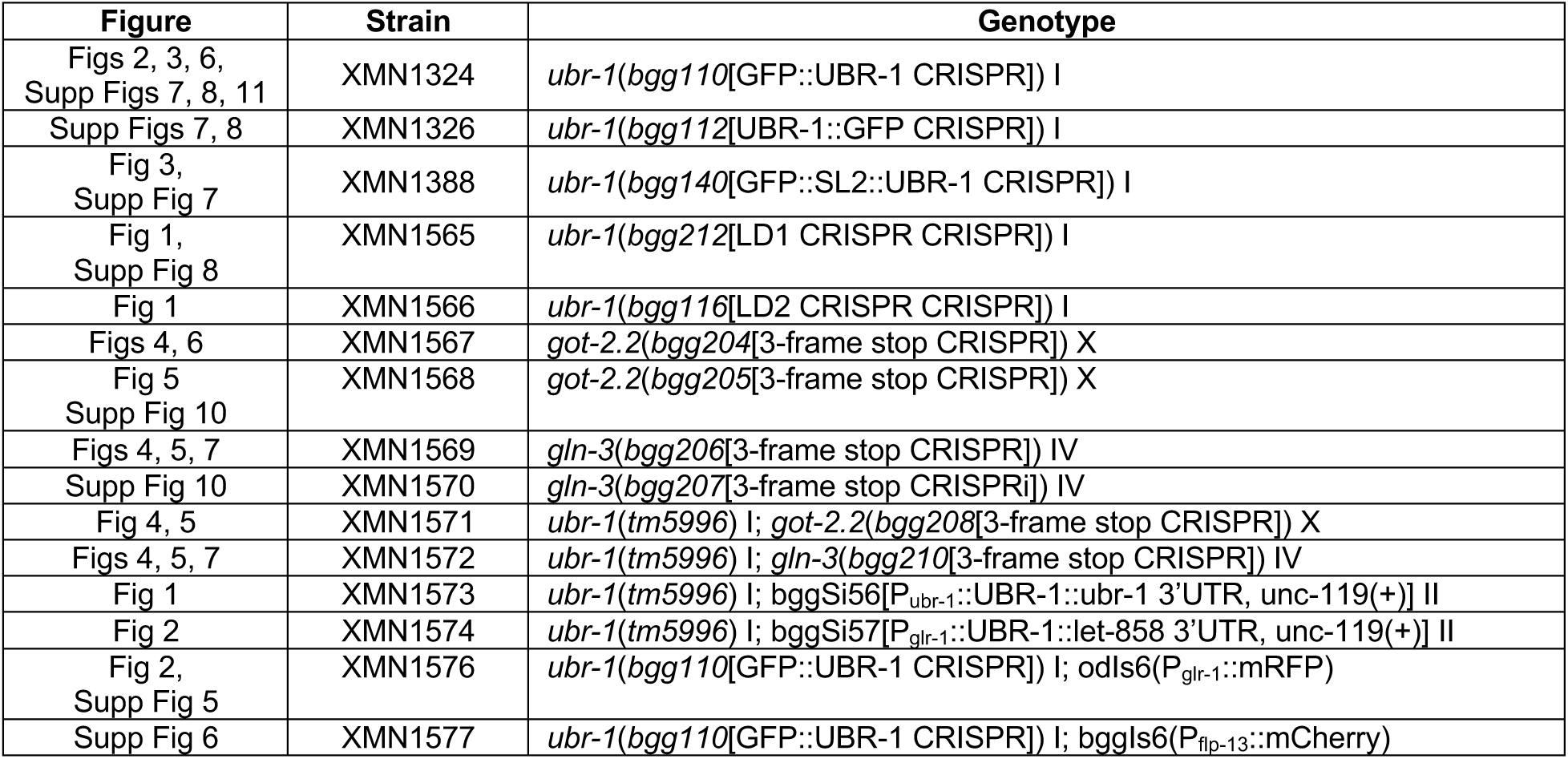
Transgenic and CRISPR Strains.

**Supplementary Table 4.**
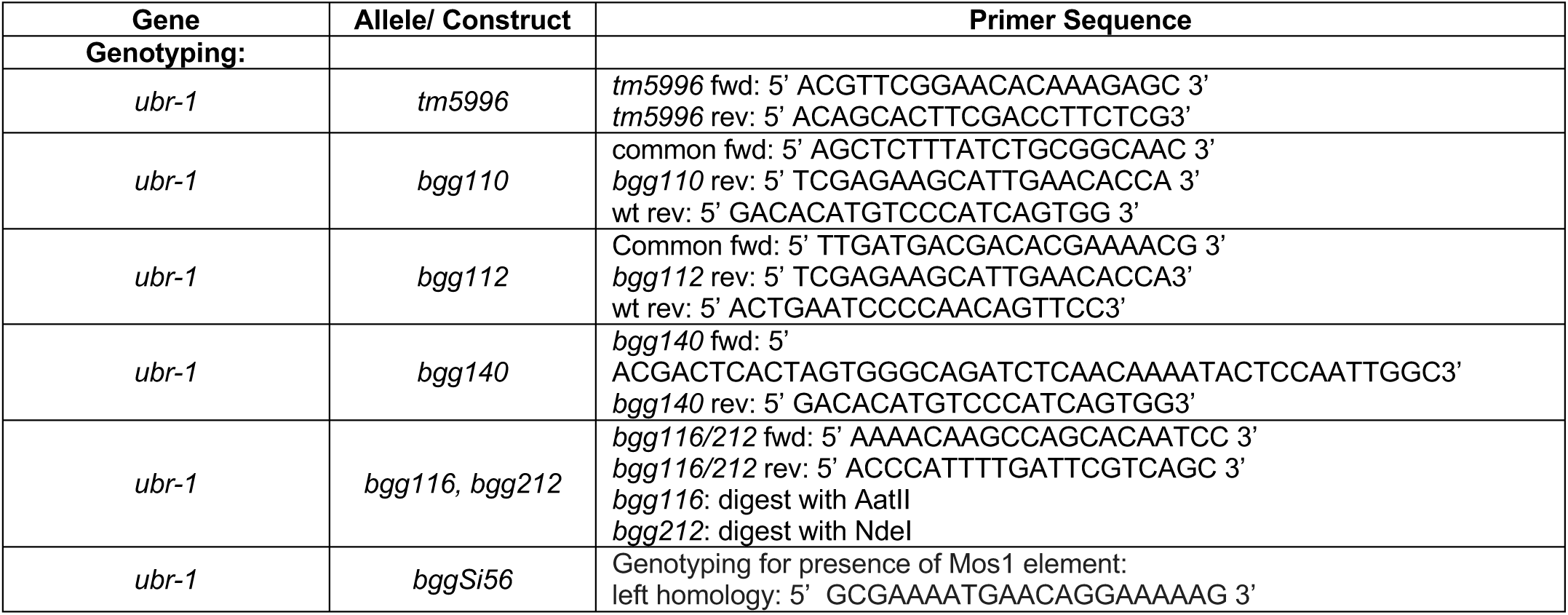
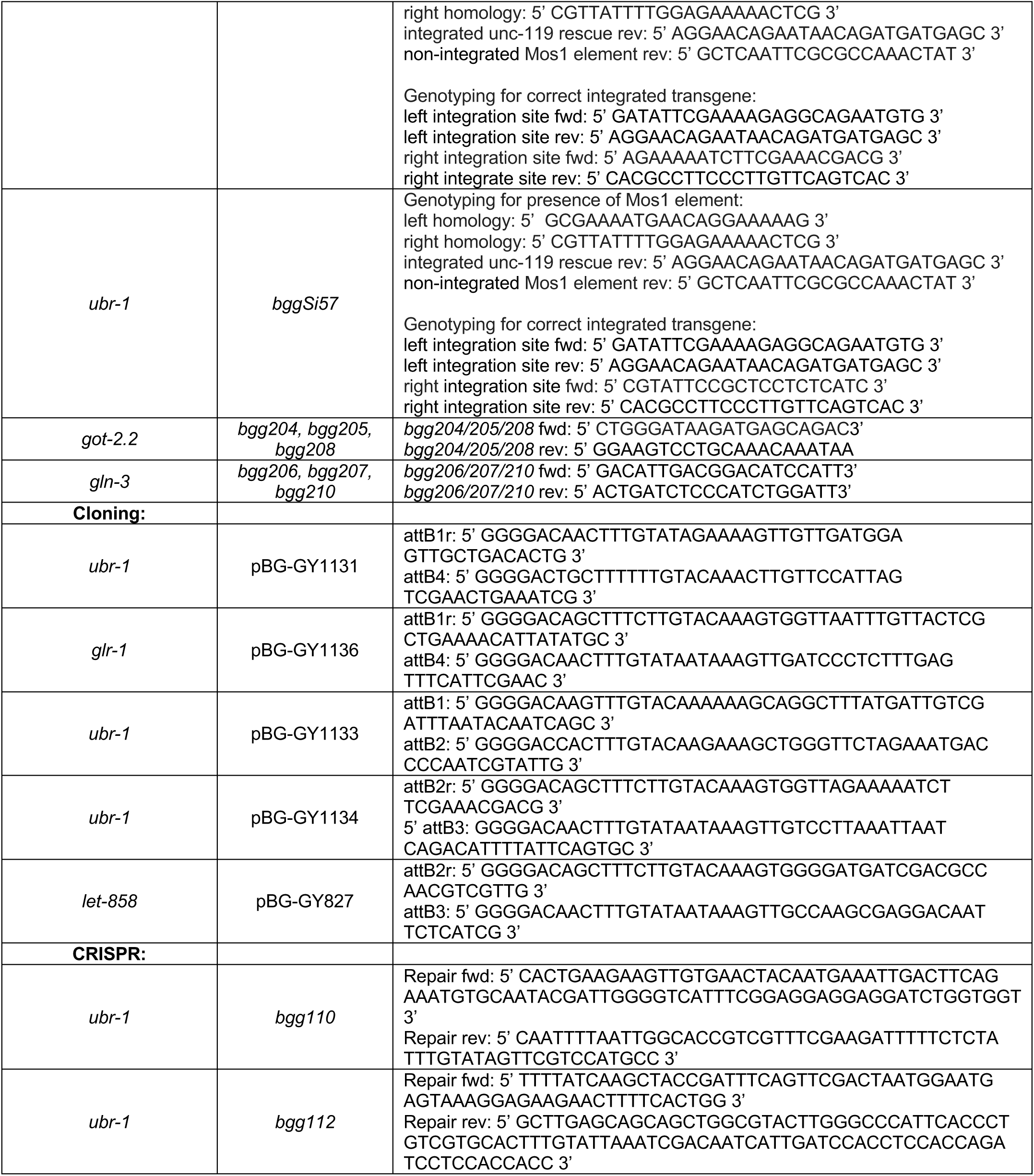
Primers.

**Supplementary Table 5.**
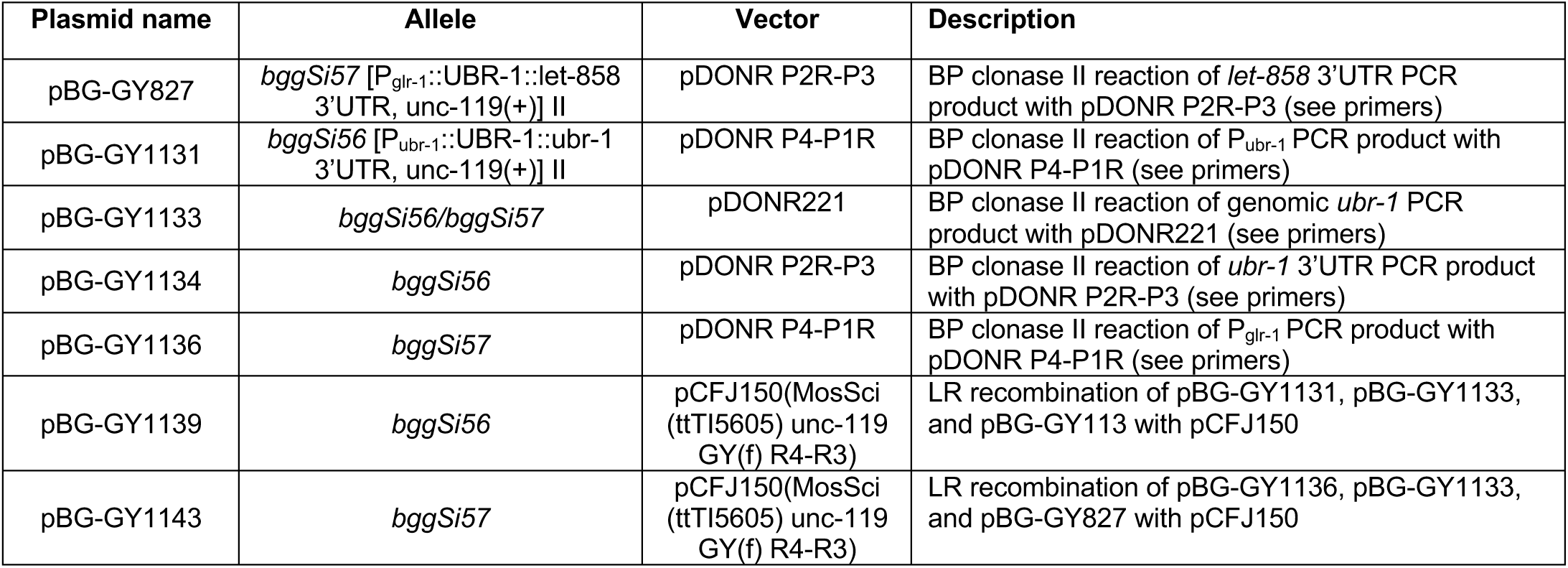
Plasmids.

**Supplementary Table 6.**
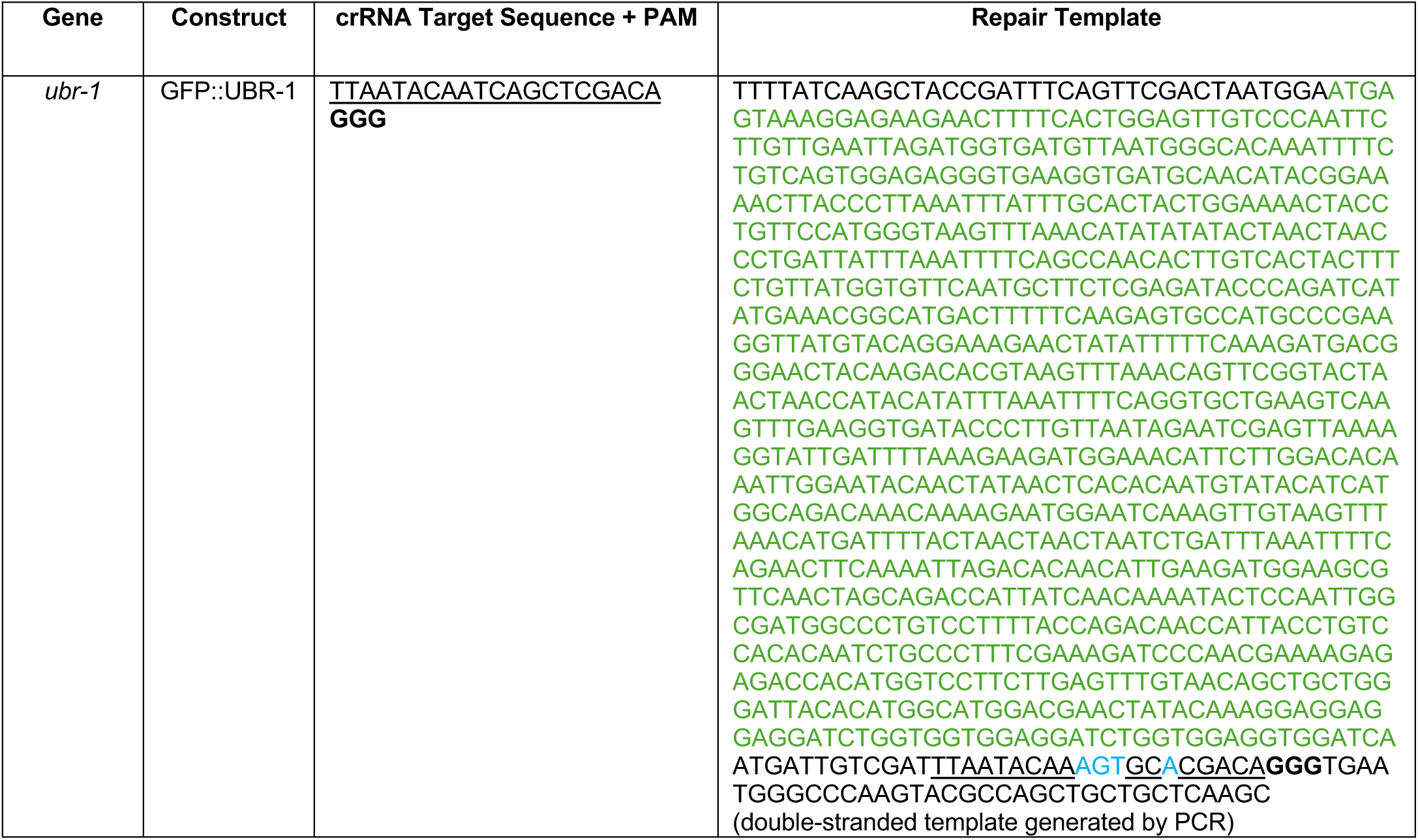
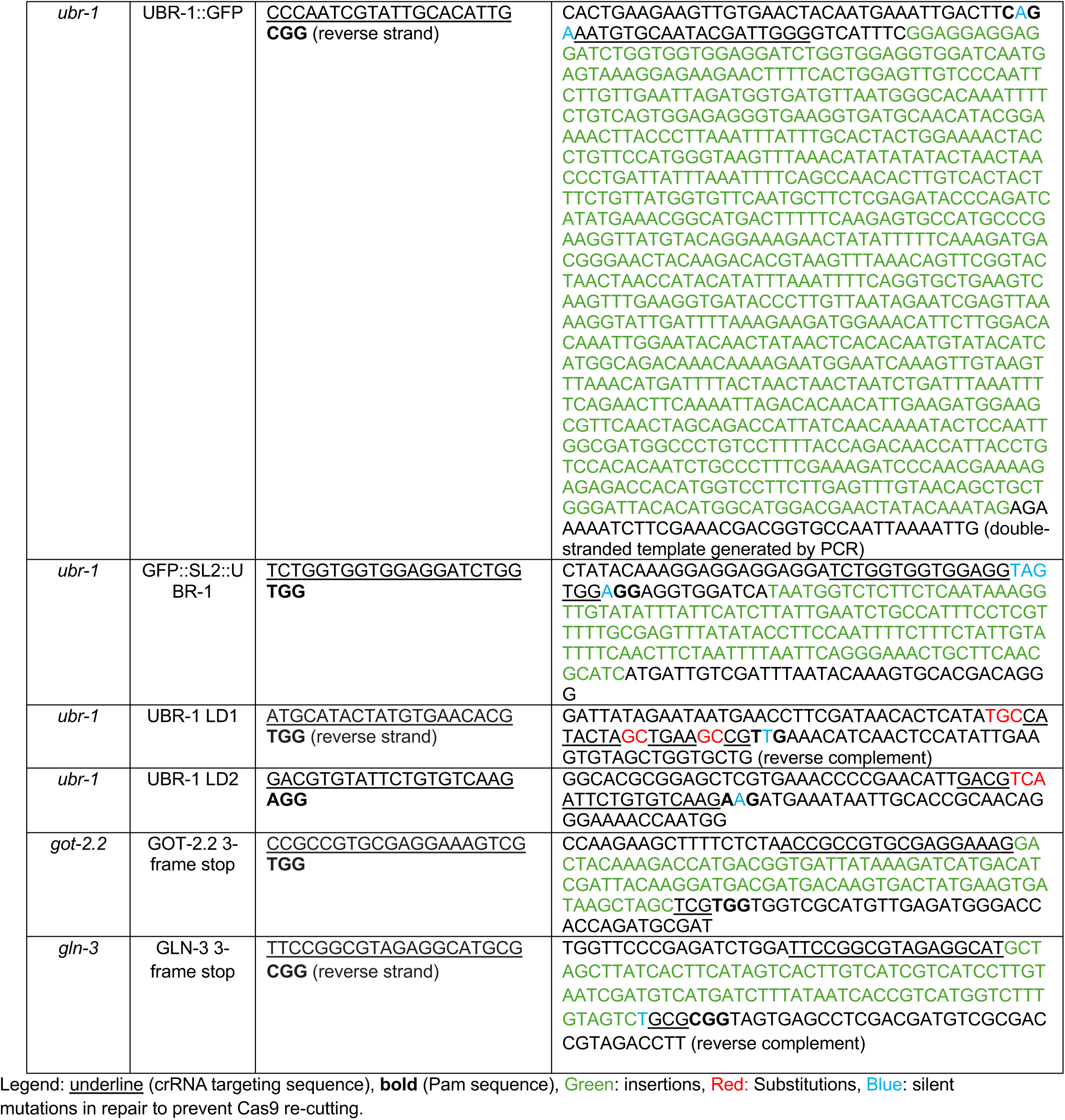
CRISPR reagents.

**Supplementary Table 7.** injection conditions.

| Figure | Transgene/Gene edit | Injected Strain | Injection Mix |
| --- | --- | --- | --- |
| Figs 2, 3, 6,<br>Supp Figs 7, 8, 11 | GFP::UBR-1 | N2 | 5.0 uL Cas9 (10 µg/µl) |
|  |  |  | 5.0 uL IDT tracrRNA (4 µg/µl) in Duplex Buffer |
|  |  |  | 0.4 ul dpy-10 crRNA (from IDT) (4 µg/µl) |
|  |  |  | 2.0 ul GFP::UBR-1 crRNA (4 µg/µl) |
|  |  |  | 5 µl GFP::UBR-1 repair oligo (4 µg) |
|  |  |  | 0.55 µl dpy-10 repair ssODN, 500 ng/µl |
|  |  |  | 2.05 µl H<sub>2</sub>O |
| Supp Figs 7, 8 | UBR-1::GFP | N2 | 5.0 uL Cas9 (10 µg/µl) |
|  |  |  | 5.0 uL IDT tracrRNA (4 µg/µl) in Duplex Buffer |
|  |  |  | 0.4 ul dpy-10 crRNA (from IDT) (4 µg/µl) |
|  |  |  | 2.0 ul UBR-1::UBR-1 crRNA (4 µg/µl) |
|  |  |  | 5 µl UBR-1::GFP repair oligo (4 µg) |
|  |  |  | 0.55 µl dpy-10 repair ssODN, 500 ng/µl |
|  |  |  | 2.05 µl H<sub>2</sub>O |
| Fig 1,<br>Supp Fig 8 | UBR-1 LD1 | N2/<br>*ubr-1(bgg110[GFP::UBR-1 CRISPR])* I | 5.0 uL Cas9 (10 µg/µl) |
|  |  |  | 5.0 uL IDT tracrRNA (4 µg/µl) in Duplex Buffer |
|  |  |  | 0.56 ul dpy-10 crRNA (from IDT) (4 µg/µl) |
|  |  |  | 2.0 ul LD1 crRNA (4 µg/µl) |
|  |  |  | 2.2 ul LD1 repair oligo (1 µg/µl) |
|  |  |  | 0.55 µl dpy-10 repair ssODN, 500 ng/µl |
|  |  |  | 4.69 µl H<sub>2</sub>O |
| Fig 1 | UBR-1 LD2 | N2 | 5.0 uL Cas9 (10 µg/µl) |
|  |  |  | 5.0 uL IDT tracrRNA (4 µg/µl) in Duplex Buffer |
|  |  |  | 0.56 ul dpy-10 crRNA (from IDT) (4 µg/µl) |
|  |  |  | 2.0 ul LD2 crRNA (4 µg/µl) |
|  |  |  | 2.2 ul LD2 repair oligo (1 µg/µl) |
|  |  |  | 0.55 µl dpy-10 repair ssODN, 500 ng/µl |
|  |  |  | 4.69 µl H<sub>2</sub>O |
| Fig 3,<br>Supp Fig 7 | GFP::SL2::UBR-1 CRISPR | *ubr-1(bgg110[GFP::UBR-1 CRISPR])* I | 5.0 uL Cas9 (10 µg/µl) |
|  |  |  | 5.0 uL IDT tracrRNA (4 µg/µl) in Duplex Buffer |
|  |  |  | 0.56 ul dpy-10 crRNA (from IDT) (4 µg/µl) |
|  |  |  | 2.0 ul LD2 crRNA (4 µg/µl) |
|  |  |  | 2.2 ul LD2 repair oligo (1 µg/µl) |
|  |  |  | 0.55 µl dpy-10 repair ssODN, 500 ng/µl |
|  |  |  | 4.69 µl H<sub>2</sub>O |
| Figs 4, 5, 6<br>Supp Fig 10 | GOT-2.2 3-frame stop CRISPR | N2/*ubr-1(tm5996)* I | 5.0 uL Cas9 (10 µg/µl) |
|  |  |  | 5.0 uL IDT tracrRNA (4 µg/µl) in Duplex Buffer |
|  |  |  | 0.56 ul dpy-10 crRNA (from IDT) (4 µg/µl) |
|  |  |  | 2.0 ul LD2 crRNA (4 µg/µl) |
|  |  |  | 2.2 ul LD2 repair oligo (1 µg/µl) |
|  |  |  | 0.55 µl dpy-10 repair ssODN, 500 ng/µl |
|  |  |  | 4.69 µl H<sub>2</sub>O |
| Figs 4, 5, 6<br>Supp Fig 10 | GLN-3 3-frame stop CRISPR | N2/*ubr-1(tm5996)* I | 5.0 uL Cas9 (10 µg/µl) |
|  |  |  | 5.0 uL IDT tracrRNA (4 µg/µl) in Duplex Buffer |
|  |  |  | 0.56 ul dpy-10 crRNA (from IDT) (4 µg/µl) |
|  |  |  | 2.0 ul LD2 crRNA (4 µg/µl) |
|  |  |  | 2.2 ul LD2 repair oligo (1 µg/µl) |
|  |  |  | 0.55 µl dpy-10 repair ssODN, 500 ng/µl |
|  |  |  | 4.69 µl H<sub>2</sub>O |
| Fig 1 | P<sub>ubr-1</sub>::UBR-1::ubr-1<br>3'UTR, unc-119(+) II | ttTi5605 II; unc-119(ed3) III; oxEx1578 | pBG-GY1139 (MosSci (unc-119) ttTi5605<br>Pubr-1(3 kb)::ubr-1 genomic::ubr-1 3'UTR)<br>10 ng/µl |
|  |  |  | pCFJ601 (Peft-3::transposase) 50 ng/µl |
|  |  |  | pMA122 (Phsp::peel-1) 10 ng/µl |
|  |  |  | pGH8 (Prab-3::mcherry) 10 ng/µl |
|  |  |  | pCFJ90 (Pmyo-2::mcherry) 2.5 ng/µl |
|  |  |  | pCFJ104 (Pmyo-3::mcherry) 5 ng/µl |
| Fig 2 | P<sub>glr-1</sub>::UBR-1::let-858<br>3'UTR, unc-119(+) II | ttTi5605 II; unc-119(ed3) III; oxEx1578 | pBG-GY1143 (MosSci (unc-119) ttTi5605<br>Pubr-1(3 kb)::ubr-1 genomic::ubr-1 3'UTR)<br>10 ng/µl |
|  |  |  | pCFJ601 (Peft-3::transposase) 50 ng/µl |
|  |  |  | pMA122 (Phsp::peel-1) 10 ng/µl |
|  |  |  | pGH8 (Prab-3::mcherry) 10 ng/µl |
|  |  |  | pCFJ90 (Pmyo-2::mcherry) 2.5 ng/µl |
|  |  |  | pCFJ104 (Pmyo-3::mcherry) 5 ng/µl |

